# *Drosophila* Clueless ribonucleoprotein particles display novel dynamics that rely on the availability of functional protein and polysome equilibrium

**DOI:** 10.1101/2024.08.21.609023

**Authors:** Hye Jin Hwang, Kelsey M. Sheard, Rachel T. Cox

## Abstract

The cytoplasm is populated with many ribonucleoprotein (RNP) particles that post-transcriptionally regulate mRNAs. These membraneless organelles assemble and disassemble in response to stress, performing functions such as sequestering stalled translation pre-initiation complexes or mRNA storage, repression and decay. *Drosophila* Clueless (Clu) is a conserved multi-domain ribonucleoprotein essential for mitochondrial function that forms dynamic particles within the cytoplasm. Unlike well-known RNP particles, stress granules and Processing bodies, Clu particles completely disassemble under nutritional or oxidative stress. However, it is poorly understood how disrupting protein synthesis affects Clu particle dynamics, especially since Clu binds mRNA and ribosomes. Here, we capitalize on *ex vivo* and *in vivo* imaging of *Drosophila* female germ cells to determine what domains of Clu are necessary for Clu particle assembly, how manipulating translation using translation inhibitors affects particle dynamics, and how Clu particle movement relates to mitochondrial association. Using Clu deletion analysis and live and fixed imaging, we identified three protein domains in Clu, which are essential for particle assembly. In addition, we demonstrated that overexpressing functional Clu disassembled particles, while overexpression of deletion constructs did not. To examine how decreasing translation affects particle dynamics, we inhibited translation in *Drosophila* germ cells using cycloheximide and puromycin. In contrast to stress granules and Processing bodies, cycloheximide treatment did not disassemble Clu particles yet puromycin treatment did. Surprisingly, cycloheximide stabilized particles in the presence of oxidative and nutritional stress. These findings demonstrate that Clu particles have novel dynamics in response to altered ribosome activity compared to stress granules and Processing bodies and support a model where they function as hubs of translation whose assembly heavily depends on the dynamic availability of polysomes.

## Introduction

Ribonucleoproteins (RNPs) often associate with cytoplasmic particles or bodies to form RNP particles, which play crucial roles in the post-transcriptional regulation of mRNAs (Gehring et al., 2017). These RNP particles, including stress granules and Processing bodies (P-bodies), are highly conserved across species and function to sequester translation machinery and mRNAs, thereby regulating mRNA stability or active translation (Anderson & Kedersha, 2006; Bauer et al., 2023; Kato & Nakamura, 2012; Keene, 2007). Under normal conditions, stress granules and P-bodies are present in limited quantities, but their numbers increase significantly in response to cellular stress (Lin et al., 2008; Schisa, 2019).

*Drosophila* Clueless (Clu) and its vertebrate analog CLUH are highly conserved multidomain ribonucleoproteins abundantly found in the cytoplasm (Fig. 1, (Cox & Spradling, 2009; Gao et al., 2014; Sen & Cox, 2016). Clu forms robust particles *in vivo*, especially in female germ cells, which exhibit high metabolic activity (Fig. 1A, B, (Cox & Spradling, 2009; Sheard et al., 2020). The loss of Clu/Cluh in *Drosophila* and mice leads to profound mitochondrial dysfunction, with flies living only a few days and mice dying on postnatal day 1 (Cox & Spradling, 2009; Schatton et al., 2017). *clu* mutants also have reduced mitochondrial proteins (Sen & Cox, 2022). Studies of CLUH have shown that a significant portion of CLUH-bound transcripts encode nucleus-encoded mitochondrial proteins, suggesting CLUH’s involvement in the regulation of mRNAs critical for mitochondrial function (Gao et al., 2014). *Drosophila* Clu also binds transcripts encoding mitochondrial proteins (Sen & Cox, 2022). The mechanism through which Clu/CLUH regulates these associated mRNAs is not yet fully understood (Hemono et al., 2022; Schatton et al., 2017; Vardi-Oknin & Arava, 2019). Moreover, Clu’s association with ribosomal proteins and translation factors suggests it plays a role in active translation, potentially involving mitochondria-associated ribosomes for co-translational or site-specific import, as well as non-mitochondria-associated cytoplasmic ribosomes (Bennett et al., 2022; Hemono et al., 2022; Pla-Martin et al., 2020; Sen & Cox, 2022; Vardi-Oknin & Arava, 2019).

**Fig 1.**
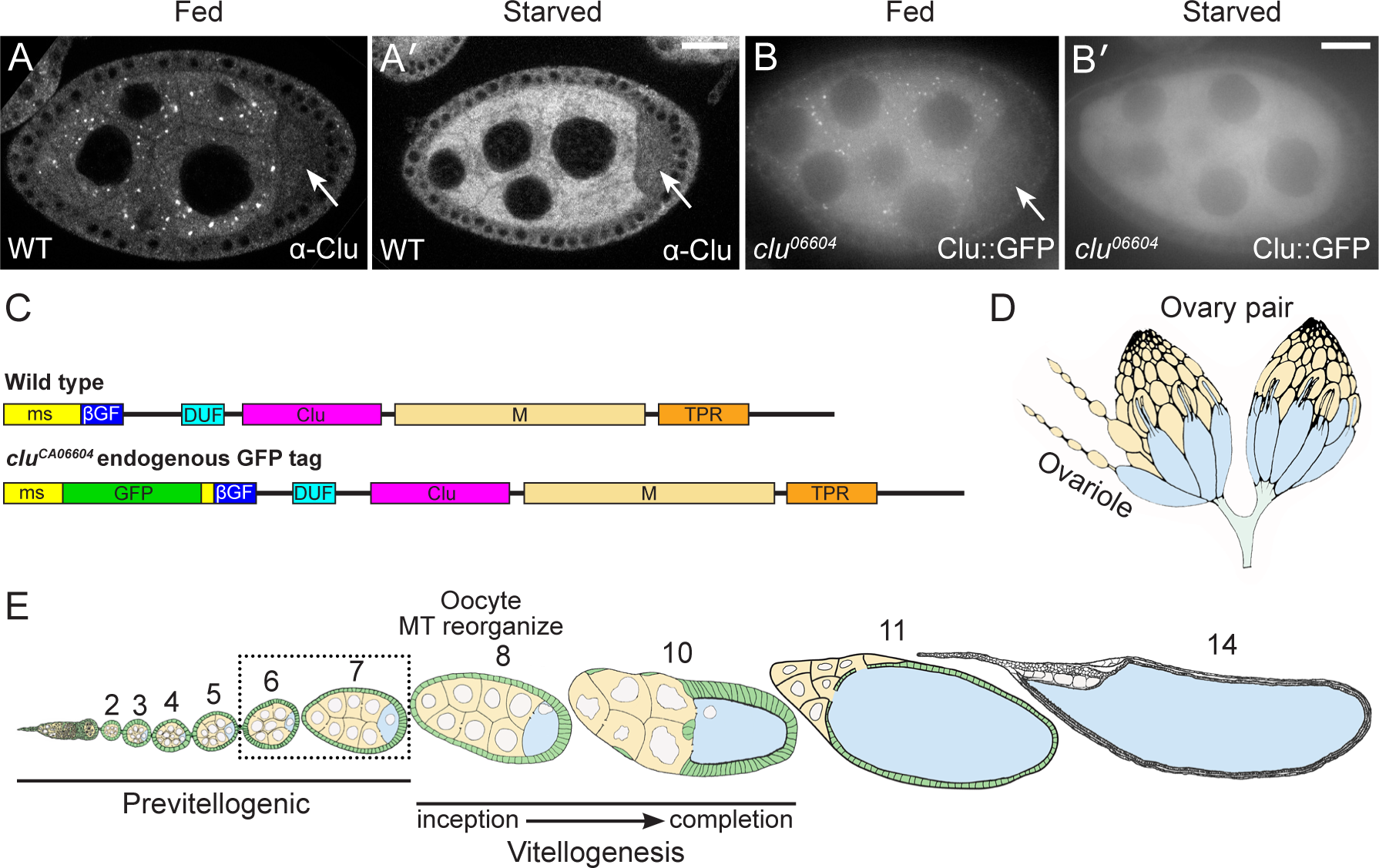
Clu forms abundant cytoplasmic particles in Drosophila nurse cells. (A-B’) Clu protein visualized in (A, A’) fixed follicles and (B, B’) still frames from live-imaged follicles that have large cytoplasmic particles in well-fed flies (A, B). Particles are disassembled with starvation (A’, B’). Clu protein is reduced in the oocyte (A, A’, B, arrows) compared to the nurse cells, and particles are absent. Fixed images were obtained using a Zeiss 700 confocal laser scanning microscope (Carl Zeiss Microscopy LLC, White Plains, NY, USA), and live images were obtained using a Nikon Eclipse Ti2 spinning disk microscope at 100x (Nikon Corporation, Tokyo, Japan). (C) Schematic showing Clu protein domains. *clu^06604^* has an in-frame Green Fluorescent Protein (GFP) inserted in the endogenous *clu* locus, resulting in a GFP fusion protein (Clu::GFP). ms = melanogaster specific, ßGF = beta Grasp Fold, DUF = Domain of unknown function, M = Middle domain, TPR = Tetratricopeptide repeat. (D, E) Schematics depicting Drosophila oogenesis. Female Drosophila have a pair of ovaries (D) composed of strings of developing follicles called ovarioles (E). (E) Ovaries from well-fed females contain all the developing follicle stages (Stages 2-14). Follicles are composed of 15 nurse cells (yellow) and one oocyte (blue) surrounded by somatic follicle cells (green). Vitellogenesis starts at stage 8 when the polarity of the oocyte’s microtubule (MT) cytoskeleton changes. Analysis and images presented in this study are predominantly stages 6 & 7 (dashed box, S1-3 Tables). (A-B’) Stage 7 follicles. White = anti-Clu antibody (A, A’) and GFP (B, B’). Scale bar = 20 µm in B’ for A-B’.

In *Drosophila*, Clu forms mitochondria-associated particles of various sizes that are highly dynamic and require intact microtubules for movement (Sheard et al., 2020). While Clu self-associates, it remains unclear whether Clu forms multimers or simply aggregates within these particles (Sen & Cox, 2016). These particles, which we have named “bliss particles,” do not co-localize with common subcellular organelle markers such as the autophagosome marker, Atg8 or the P-body protein, Trailer Hitch (Sheard et al., 2020). Unlike stress granules and P-bodies, bliss particles are exquisitely sensitive to stress *in vivo* and only form under optimal conditions in well-fed flies (Sheard et al., 2020). Particle disassembly can be induced by starvation, oxidative stress, and mitochondrial stress caused by mutations in *Superoxide Dismutase 2* (*Sod2*), *PTEN-induced putative kinase 1* (*Pink1*), and *parkin* (Fig. 1A’, B’, (Cox & Spradling, 2009; Sen et al., 2015; Sheard et al., 2020). Remarkably, removing stress conditions, such as by refeeding the flies or by adding insulin *ex vivo*, restores bliss particles (Sheard et al., 2020). Despite the dynamic nature of these particles, Clu levels remain constant, indicating that particle disassembly is not due to protein degradation (Sheard et al., 2020). Furthermore, insulin signaling is necessary and sufficient for particle assembly, suggesting that the cell’s metabolic state significantly influences the presence of bliss particles (Sheard et al., 2020).

Mitochondria, as central hubs of metabolism, house critical pathways such as heme biosynthesis, fatty acid ß-oxidation, and steroidogenesis, in addition to generating ATP (Bennett et al., 2022; Rahman, 2020; Scarpulla, 2008). While these organelles have their own mitochondrial DNA, the majority of proteins required for these pathways are supplied by nuclear DNA-encoded mRNAs that are translated on cytoplasmic and mitochondrial-associated ribosomes (Bennett et al., 2022; Devaux et al., 2010; Dimmer et al., 2002). Given Clu’s binding to mRNAs and its association with ribosomes, as well as the juxtaposition of bliss particles to mitochondria, this study aims to determine how disrupting translation affects bliss particle dynamics in *Drosophila* female germ cells.

We employed ectopic transgenic expression *in vivo* to demonstrate that three conserved domains of Clu are necessary for its assembly into particles, and that an excess of full-length Clu disassembles particles, suggesting particle stability or assembly is regulated by cytoplasmic Clu concentration. Manipulating translation acitivity using the translation inhibitors puromycin (PUR) and cycloheximide (CHX) revealed that PUR treatment quickly disassembled Clu bliss particles whereas CHX treatment did not. In addition, CHX treatment stabilized existing bliss particles in the presence of nutritional and oxidative stress. While CHX treatment did not interfere with directed movement of bliss particles, by analyzing size and speed of Clu particles associated with mitochondria, we found that particles bound to mitochondria move more slowly. Together, these observations support a model whereby *Drosophila* Clu bliss particle assembly requires polysomes and that they could function as sites of active translation of mRNAs encoding mitochondrial proteins to regulate mitochondrial function. In addition, our observed particle dynamics are distinct and unique from those observed for other RNP particles, underscoring the distinctive response of Clu bliss particles to translation inhibition in *Drosophila* germ cells.

## Results

### *Drosophila* ovaries as a model to study Clu bliss particle dynamics

To identify the protein domains and protein synthesis requirements of Clu particle dynamics, we examined particle assembly and disassembly in female germ cells using fixed and live imaging. Clu particles are always present in the nurse cells of well-fed follicles (Cox & Spradling, 2009; Sheard et al., 2020). However, they look larger or smaller, or more or less numerous, depending on imaging conditions, including the length of dissection, whether the tissue is imaged while fixed or live, and the type of microscope (spinning disc vs laser confocal). However, importantly, Clu particles completely disassemble in response to stress (Fig 1A’, B’), thus we used a binary decision for assembly/disassembly – either Clu particles were present or absent – to determine how various conditions affect dynamics. *Drosophila* females have a pair of ovaries composed of 16-20 ovarioles (Fig 1D, E, (Spradling, 1993) (Sarikaya et al., 2012)). Ovarioles are strings of developing follicles that contain all follicle stages (2-14) when dissected from well-fed females. For consistency, we predominantly analyzed data from Stage 6 & 7 follicles, which are large enough to be readily imaged and contain many Clu particles but are previtellogenic, occurring before many complex developmental events (Fig 1E, S1 Table, see Materials and Methods for additional details regarding imaging).

### The DUF, Clu, and TPR domains are necessary for Clu bliss particle association

Clu is a large, multi-domain protein (Fig 1C); however, the function of each domain and how they control particle dynamics is poorly understood. The ms (melanogaster-specific) domain is unique to *Drosophila* and is not required to rescue the *clu* null mutant (Sen & Cox, 2016). The ßGF (beta Grasp Fold) domain is so-called based on the predicted structure, and the DUF (Domain of Unknown Function) is predicted based on sequence homology (Sen et al., 2015). The Clu domain is highly conserved, but we do not yet understand its role. We previously demonstrated that the TPR (tetratricopeptide repeat) domain is critical for Clu’s binding mRNA (Sen & Cox, 2016). Finally, the M (Middle) domain sequence is unstructured. While it does not contain traditional intrinsically disordered (ID) motifs, ID motifs are characteristic of proteins associated with membraneless organelles (Alberti et al., 2019; Li et al., 2012; Shin & Brangwynne, 2017).

Previously, we used co-immunoprecipitation to show that full-length *Drosophila* Clu can self-associate (Sen & Cox, 2016). To test which domains are required for Clu particle assembly *in vivo*, we used the GAL4/UAS system. We created transgenic lines that ectopically express Clu-tagged with mScarlet at the C-terminus under the control of the conditional UASp promoter (Fig 2A). To simultaneously visualize ectopically expressed mScarlet-labeled constructs and endogenous Clu, each construct was combined with *nanos* GAL4 (*nos*GAL4) in a *clu^06604^*background which expresses Clu::GFP (Fig 1C). The *nos*GAL4 line we used is clonally expressed in the nurse cells (Hanyu-Nakamura et al., 2004). Using live imaging, we found full-length (FL)-Clu::mScarlet reliably formed Clu particles (Fig 2B’), which co-localized with endogenous Clu::GFP particles, indicating that both Clu species exist within the same particle (Fig 2B”, S1 Movie). To determine if the DUF, Clu, and TPR domains are required for particle assembly, we ectopically overexpressed mScarlet-labeled Clu transgenes with each domain deleted (ΔDUF::mScarlet, ΔClu::mScarlet, and ΔTPR::mScarlet) (Fig 2A, C-E”, S2-4 Movies). Using live imaging, we were unable to see particle assembly of any of these deletion constructs (Fig 2C’, D’, E’). Furthermore, none of these deletions co-labeled with endogenous Clu::GFP particles (Fig 2C”, D”, E”). This suggests that these three domains are necessary to assemble Clu particles and to associate with already assembled endogenous particles.

**Fig 2.**
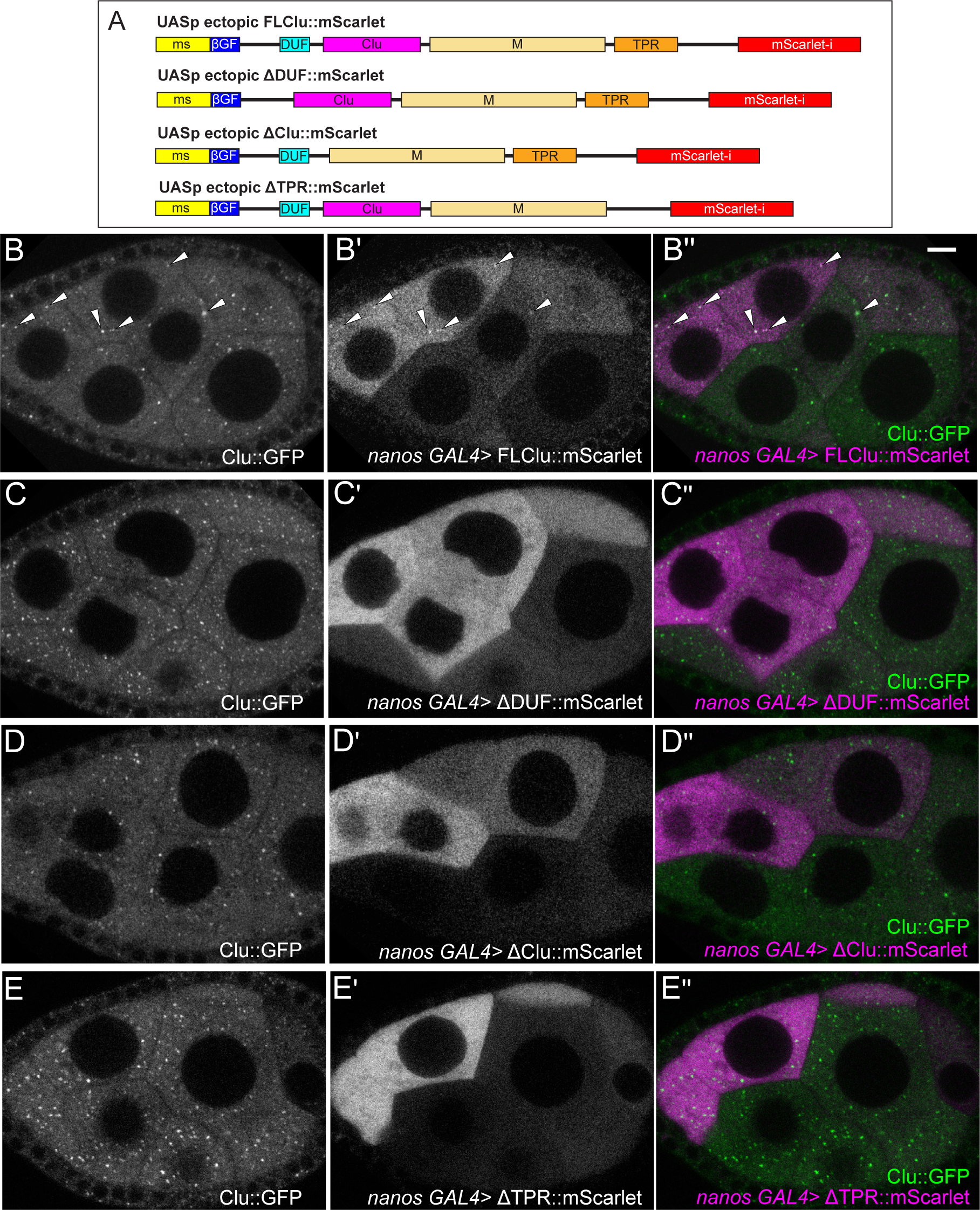
DUF, Clu, and TPR domains are required for Clu particle association. (A) Cartoon of full-length (FL) and domain deletion (ΔDUF, ΔClu, and ΔTPR) constructs of ectopic Clu tagged with the fluorescence tag mScarlet. (B-B”) Still-images from S1 Movie of a follicle from a *clu^CA06604^* /*+*; *nanos (nos) GAL4*/*UASp-FLclu*::*mScarlet* female. Particles from endogenous Clu::GFP (B) and ectopic FLClu::mScarlet (B’) co-localize in germ cells (B”) as arrow heads indicate. 71% of nurse cells expressing mScarlet showed colocalization of Clu::GFP and mScarlet (n=14 follicles, see S1 Table for details). (C-E”) Still-images from ectopic Clu deletion constructs (S2-4 Movies). Endogenous Clu::GFP (C, D, E) forms particles. However, ΔDUF (C’), ΔClu (D’), and ΔTPR (E’) constructs do not form particles (C’, D’, E’) and cannot associate with endogenous Clu particles (C”, D”, E”, See S1 Table for details). Still-images from (C-C”) S2 Movie of a follicle from *clu^CA06604^*/+*; nosGAL4*/*UASp-clu*Δ*DUF*::*mScarlet,* (D-D’’) S3 Movie of a follicle from *clu^CA06604^*/+*; nosGAL4*/*UASp-clu*Δ*Clu*::*mScarlet*, and (E-E’’) S4 Movie of a follicle from *clu^CA06604^*/+; *nosGAL4*/*UASp-clu*Δ*TPR*::*mScarlet* females. (B-E”) Stage 7 egg chamber follicles expressing Clu::GFP and various ectopic Clu tagged with mScarlet were imaged with a 200 µg/mL insulin-containing Complete Schneider’s (CS) media in a time-lapse course at a single plane (see S1-4 Movies for details). The focal plane was selected by ensuring more than three nurse cells having nuclear and cytosolic area clearly were visible, with approximately 25 % depth from the top surface of each follicle (See Materials and Methods for details). Live images were obtained using a Nikon A1 plus Piezo Z Drive Confocal microscope at 60x (Nikon Corporation, Tokyo, Japan). The follicle stages analyzed (n) in each genotype: *clu^CA06604^*/*+*; *nanos (nos) GAL4*/*UASp-FLclu*::*mScarlet*, stage 5 (2), stage 6 (3), stage 7 (8), stage 8 (1); *clu^CA06604^*/+*; nosGAL4*/*UASp-clu*Δ*DUF*::*mScarlet*, stage 6 (2), stage 7 (4), stage 8 (2); *clu^CA06604^*/+*; nosGAL4*/*UASp-clu*Δ*Clu*::*mScarlet*, stage 5 (1), stage 6 (4), stage 7 (5), stage 8 (1); *clu^CA06604^*/+;*nosGAL4*/*UASp-clu*Δ*TPR*::*mScarlet*, stage 6 (3), stage 7 (5), stage 8 (3). More details, including the number of nurse cells expressing mScarlet and having Clu particles, the number of nurse cells showing colocalization of endogenous GFP and ectopic mScarlet, and the number of dissected animals, are described in S1 Table. (B, C, D, E) White = endogenous Clu::GFP. (B’, C’, D’, E’) White = mScarlet. (B”, C”, D”, E”, merge) Green = Clu::GFP, magenta = Scarlet. Scale bar = 10 µm in B” for B-E”.

### Bliss particles disassemble in response to high expression of functional Clu

When we tested expression levels of ectopic FL-Clu using different GAL4 drivers, we noticed a dose-dependent effect on particle formation. *daughterless* (*da)* GAL4 induced high uniform expression of Clu in germ cells (Fig 3A-A”) (Casper et al., 2011; Morawe et al., 2011). In contrast to *nos*GAL4, when ectopic FL-Clu::mScarlet was expressed by *da*GAL4 (Fig 3A’), neither ectopic Clu nor endogenous Clu assembled into particles (Fig 3A vs 2B-B”). To test whether this was dependent on functional FL-Clu, we ectopically overexpressed Scarlet-labeled ΔDUF or ΔTPR domain (ΔDUF::mScarlet, ΔTPR::mScarlet, Fig 2A) using *da*GAL4. Consistent with our observations using *nos*GAL4, neither was able to assemble mScarlet particles (Fig 3B’, C’), but unlike ectopic FL-Clu, ΔDUF and ΔTPR overexpression did not disassemble endogenous Clu particles (Fig 3B, C). Finally, to ensure that expressing high concentrations of any protein does not cause cell stress that inhibits particle assembly, we overexpressed unrelated mCherry-labeled Capping Protein Beta (mCherry::CPB) (Ogienko et al., 2013). mCherry::CPB also did not disrupt endogenous Clu particles (Fig 3D-D”). This suggests that the mechanism assembling Clu particles at least partly depends on regulating the concentration of functional Clu but does not respond to non-functional Clu (ΔDUF and ΔTPR).

**Fig 3.**
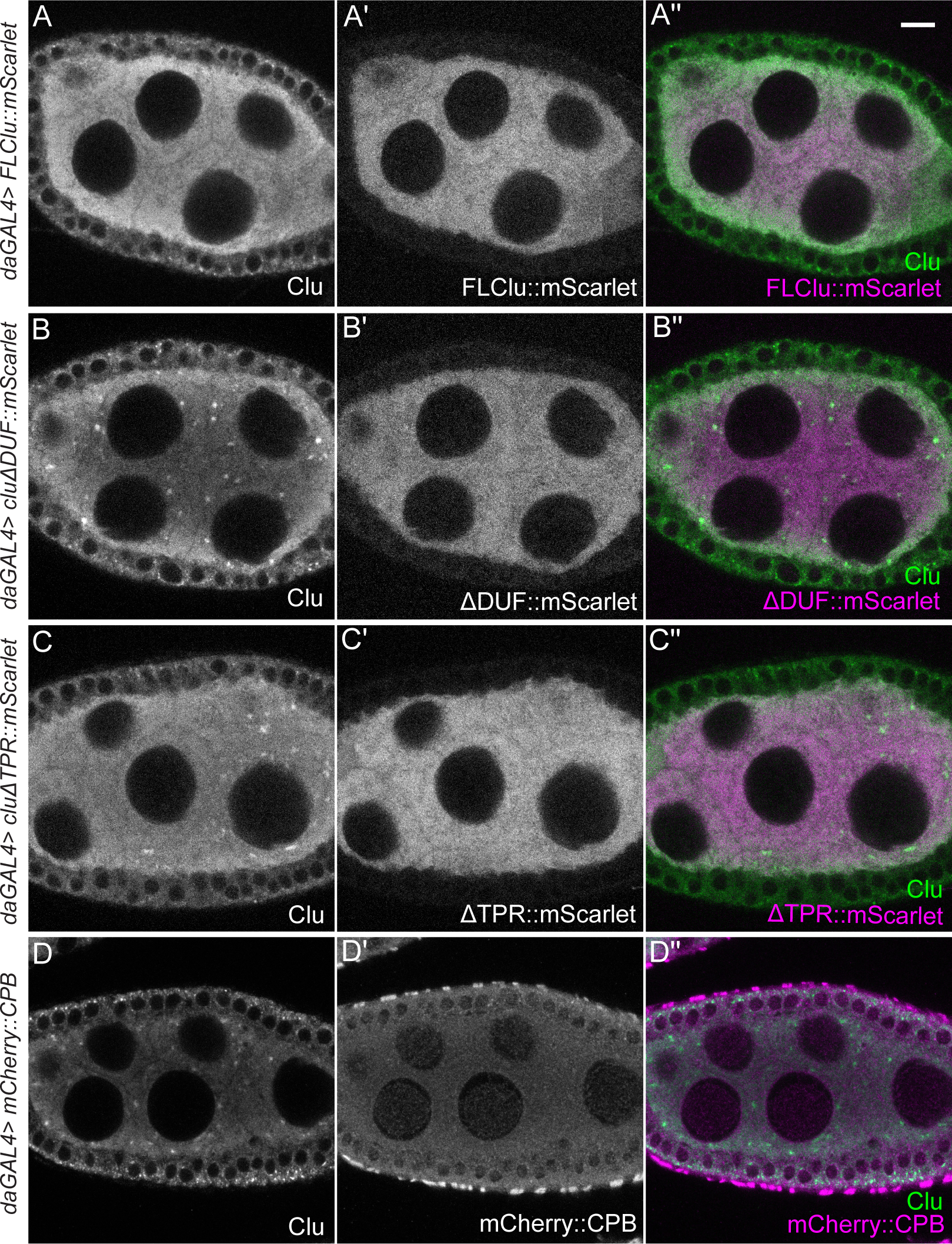
High levels of functional Clu disassemble bliss particles. (A-A”) Immunostaining of a follicle from a *daughterless (da) GAL4*/*UASp-FLclu*::*mScarlet* female. High levels of ectopic FLClu (A’) disrupt particle formation (A, A”). (B-B”) Immunostaining of a follicle from a *daGAL4*/*UASp-clu*Δ*DUF*::*mScarlet* female. (C-C”) Immunostaining of a follicle from a *daGAL4*/*UASp-clu*Δ*TPR*::*mScarlet* female. (D-D”) Immunostaining of a follicle from*daGAL4*/*UASp-mCherry*::*cpb* female. High levels of ΔDUF (B’), ΔTPR (C’) or CPB (D’) do not interfere with endogenous Clu particle formation (B, B” for ΔDUF, C, C” for ΔTPR or D, D” for CPB). 95% (ΔDUF), 90% (ΔTPR), and 94% (CPB) of nurse cells expressing each ectopic construct from the observed follicles showed Clu particles (See S1 Table for details). (A-D”) Stage 7 egg chamber follicles were imaged with a 1.2 µm thickness of z-stacks with an interval of 0.42 µm. The focal plane was selected by ensuring at least three to four nuclei were clearly visible in nurse cells, but also to avoid dim fluorescence signals due to deeper depth (See Materials and Methods for details). Images were obtained using a Zeiss LSM 980 confocal laser scanning microscope (Carl Zeiss Microscopy LLC, White Plains, NY, USA). The follicle stages analyzed (n) for each genotype by immunostaining: *daGAL4*/*UASp-FLclu*::*mScarlet*, stage 5 (1), stage 6 (3), stage 7 (4), stage 8 (1); *daGAL4*/*UASp-clu*Δ*DUF*::*mScarlet*, stage 5 (1), stage 6 (1), stage 7 (1), stage 8 (1); *daGAL4*/*UASp-clu*Δ*TPR*::*mScarlet*, stage 5 (1), stage 6 (1), stage 7 (2), stage8 (1); *daGAL4*/*UASp-mCherry*::cpb, stage5 (1), stage 6 (2), stage 7 (4), stage 8 (4). More details, including the number of nurse cells having Clu particles and the number of dissected animals, are described in S1 Table. (A, B, C, D) White = anti-Clu, (A’, B’, C’) white = anti-Scarlet, (D’) white = anti-mCherry. (A”, B”, C”, merge) Green = anti-Clu, magenta = anti-Scarlet. (D”, merge) Green = anti-Clu, magenta = anti-mCherry. For A, A”, B, B”, C, C”: Note: anti-Clu antibody also recognizes the mScarlet transgene. Scale bar = 10 µm in A” for A-D”.

### The translation inhibitor puromycin disassembles Clu bliss particles

Stress granules and P-bodies have distinct responses of assembly/disassembly in response to various stressors and different translation inhibitors, depending on the drugs’ mechanism (Buddika et al., 2020; Eulalio et al., 2007; Patel et al., 2016). This is thought to be due to manipulation of available levels of associated messenger RNPs (mRNPs) and polysome presence and activity (Hofmann et al., 2021; Hubstenberger et al., 2017; Ivanov et al., 2019; Kato & Nakamura, 2012). Puromycin (PUR) is a commonly used and well-studied translation inhibitor. PUR forms a stable peptide bond with nascent polypeptides, resulting in premature translation termination which leads to ribosome complex disassembly, polysome loss and increased free mRNAs (Aviner, 2020) (Fig. 4A). Since Clu is a ribonucleoprotein critical for mitochondrial function that binds mRNAs encoding mitochondrial proteins, we wanted to determine how puromycin treatment affected bliss particle dynamics. To do this, we treated ovarioles dissected from well-fed *clu^06604^* females with PUR and found that the plentiful bliss particles were quickly and completely disassembled within ten minutes (Fig 4B-C’, S5 Movie). Given the molecular action of PUR, this suggests that disassembled ribosomes and/or increased concentrations of mRNPs cause Clu particles to disassemble.

**Fig 4.**
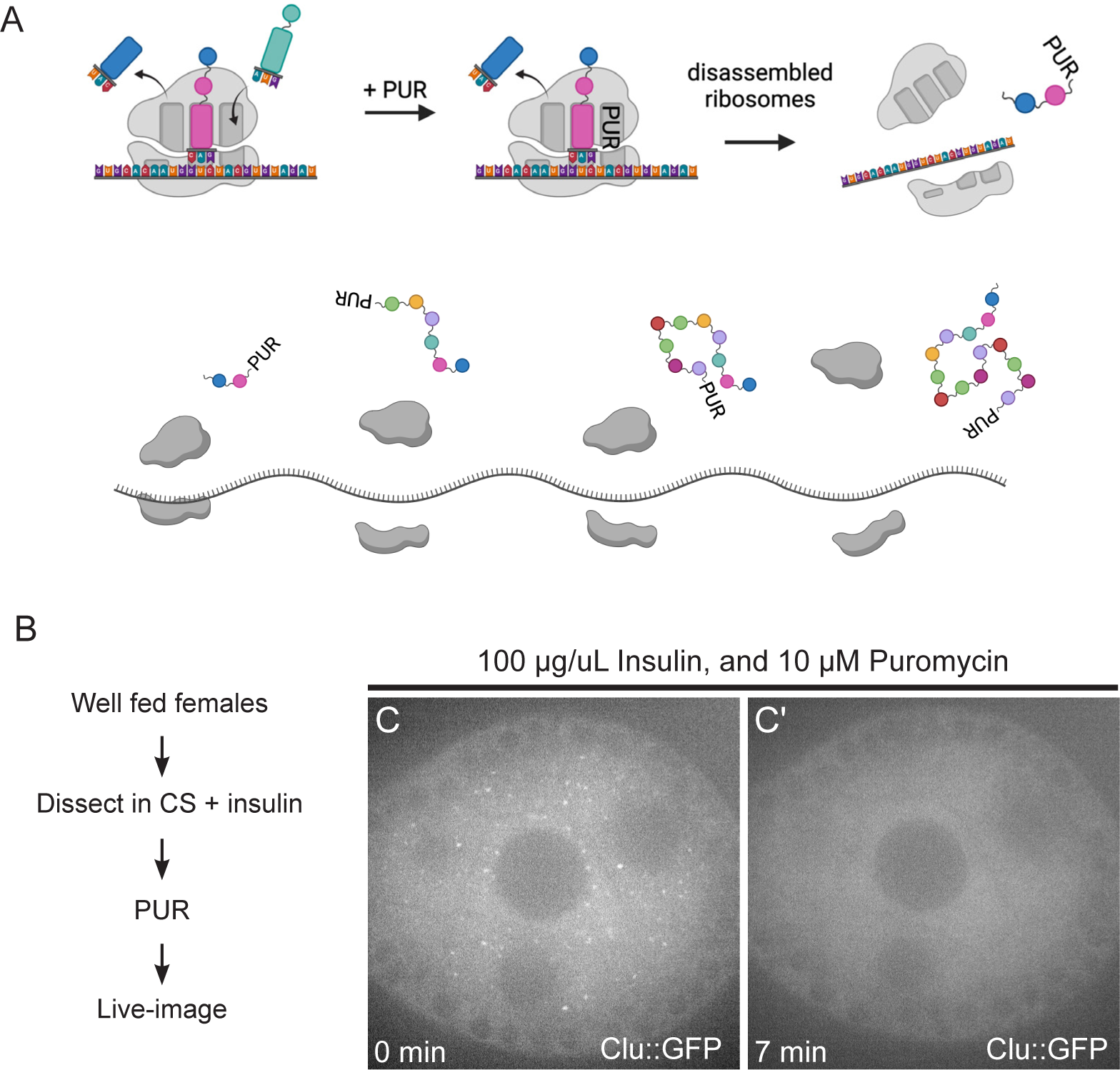
The translation inhibitor puromycin disassembles Clu bliss particles. (A) Schematic demonstrating the mechanism of action for the translation inhibitor, puromycin (PUR). PUR blocks nascent polypeptide chain elongation, thereby causing premature translation termination, disassembly of the ribosomal complex, and decreased polysomes. (B) Workflow for the experiment. Ovarioles dissected from Well-fed *clu^CA06604^* females were treated with puromycin, then live-imaged (S5 Movie, C, C’). (C) The 1^st^ still frame (at time zero after adding 10 µM PUR) of stage 6 follicle from S5 Movie showing Clu particles. (C’) The 22^nd^ still-image (at seven minutes) of the same follicle demonstrating disassembled bliss particles by PUR (n=20/20 follicles, see S2 Table for details). The focal plane was selected by ensuring at least three to four nuclei were clearly visible in nurse cells, with approximately 25 % depth from the top surface of each follicle (See Materials and Methods for details). Live images were obtained using a Nikon Eclipse Ti2 spinning disk microscope at 100x (Nikon Corporation, Tokyo, Japan). The follicle stages analyzed (n): stage 5 (2), stage 6 (8), stage 7 (5), stage 8(5). Scale bar = 10 µm in C’ for C and C’.

### Cycloheximide treatment maintains Clu bliss particles, but blocks insulin-induced assembly

CHX is another well-known and frequently used translation inhibitor that binds to the exit site of the ribosome which stalls the ribosome and blocks translation elongation, leading to increased concentrations of polysomes (Fig 5A) (Duncan & Mata, 2017).

**Fig 5.**
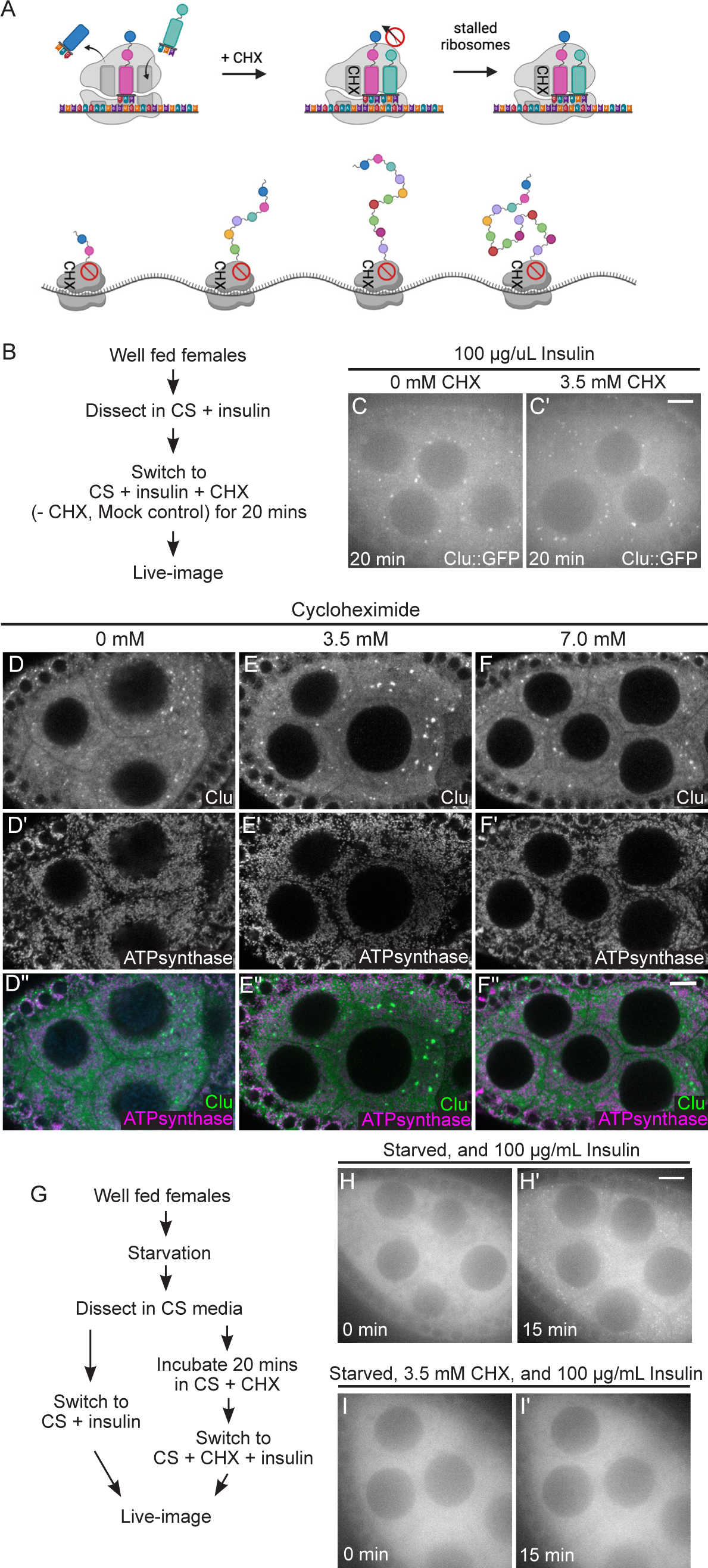
The translation inhibitor cycloheximide maintains Clu bliss particles, but blocks insulin-induced assembly. (A) Schematic demonstrating the mechanism of action for the translation inhibitor, cycloheximide (CHX). CHX blocks the 60S ribosome exit site, thereby stalling translation and increasing polysome densities. (B) Workflow for the CHX treatment showing in C and C’. Ovarioles from well-fed *clu^CA06604^* females were incubated with CHX for 20 minutes then live-imaged (S6 and 7 Movies, C, C’). (C) Still-image of a stage 7 follicle after 20-minute incubation without CHX demonstrating the presence of Clu bliss particles (n=14/14 follicles, see S2 Table for details). (C’) Still-image of a stage 7 follicle after 20-minute incubation with 3.5 mM CHX demonstrating the assembled bliss particles are still present (n=20/20 follicles, see S2 Table for details). Follicles were imaged in a time-lapse course for 3 minutes at a single plane at the end of 20 minutes. The focal plane was selected by ensuring at least three to four nuclei were clearly visible in the nurse cells, with approximately 25 % depth from the top surface of each follicle (See Materials and Methods for details). Follicle stages analyzed (n): mock treatment, stage 5 (3), stage 6 (3), stage 7 (7), stage 8 (1); 3.5 mM CHX treatment, stage 5 (3) stage 6 (4), stage 7 (10), stage 8 (3). (D-F”) Immunostaining of stage 7 follicles from well-fed *w^1118^* females fed for 24 hours with wet yeast paste containing (D-D”) 0 mM, (E-E”) 3.5 mM, and (F-F”) 7 mM CHX. None of CHX feeding disassemble bliss particles (D, E, F) and normal mitochondrial morphology and localization is maintained (D’, E’, F’). Observed bliss particles: 91% (0 mM CHX), 94% (3.5 mM CHX), and 96% (7 mM CHX) (See S2 Table for details) of nurse cells showed. Images are 2 µm projections assembled 0.42 µm sections. The focal plane was selected to show at least 3∼4 nuclei but also to avoid dim fluorescence signals due to deeper depth (See Methods for details). The total number of follicles analyzed (n) by immunostaining: 0 mM CHX, stage 6 (1), stage 7 (5), stage 8 (6); 3.5 mM CHX, stage 5 (6), stage 6 (5), stage 7 (6), stage 8 (4); 7 mM CHX, stage (7), stage 6 (6), stage 7 (10), stage 8 (6). (G) Workflow for the CHX experiment showing in (H-I’). Well-fed *clu^CA06604^* females were starved for 3 hours with water only, then ovarioles dissected from starved animals were incubated in insulin (H, H’, control) or CHX followed by insulin (I, I’). (H) The 1^st^ still frame (at time zero after adding 100 µg/mL insulin) of stage 7 follicle from S8 Movie showing no Clu particles. (H’) The 46^th^ still-image (at 15 minutes) of the same follicle from S5 Movie demonstrating the recovery of bliss particles by insulin as previously showed (Sheard, 2020) (n=4/4 follicles, see S2 Table for details). (I) The 1^st^ still frame (at time zero after adding 100 µg/mL insulin) of stage 7 follicle starved and treated with CHX from S9 Movie. (I’) The 46^th^ still frame (at 15 minutes) of the same follicle showing no recovery of bliss particle by insulin following CHX treatment (n=11/11 follicles, see S2 Table for details). The focal plane was selected by ensuring at least three to four nuclei were clearly visible in the nurse cells, with approximately 25 % depth from the top surface of each follicle (See Materials and Methods for details). Follicles stages analyzed (n): insulin only treated, stage 5 (1), stage 7 (2), stage 8 (1); insulin following 3.5 mM CHX, stage 5 (3) stage 6 (4), stage 7 (4). More details, including the number of nurse cells having Clu particles and the number of dissected animals, are described in S2 Table. Live images were obtained using a Nikon Eclipse Ti2 spinning disk microscope at 100x (Nikon Corporation, Tokyo, Japan). Immunostaining images were obtained using a Zeiss LSM 980 confocal laser scanning microscope (Carl Zeiss Microscopy LLC, White Plains, NY, USA). (D, E, F) White = anti-Clu. (D’, E’, F’) White = anti-ATP synthase. (D”, E”, F”, merge) Green = anti-Clu, magenta = anti-ATP synthase. Scale bar = 10 µm in B’ for B-C’, in F” for D-F”, and in H’ for H-I’.

Under normal conditions, CHX treatment causes P-bodies to disassemble (Eulalio et al., 2007; Kedersha et al., 2000; Lin et al., 2008; Moutaoufik et al., 2014; Patel et al., 2016). To determine the effect of CHX on bliss particles *ex vivo*, we tested whether CHX treatment disassembled particles and found that this was not the case (Fig. 5B-C’). To ensure our method of CHX treatment was effective, we dissected ovarioles from well-fed *trailer hitch^CA06517^* (*tral*) females that express GFP inserted at the endogenous *tral* locus, thus labeling P-bodies (Buszczak et al., 2007; Eulalio et al., 2007; Kato & Nakamura, 2012). We confirmed that Tral-labeled P-bodies decreased in size and number with CHX addition compared to mock control, as previously shown (S1 Fig) (Eulalio et al., 2007; Patel et al., 2016).

*Ex vivo* imaging has many advantages for investigating cellular dynamics. However, we wanted to determine the effect of CHX on Clu bliss particles *in vivo*. To do this, we switched well-fed females to yeast paste supplemented with CHX for 24 hours. Feeding CHX has been shown to reduce protein synthesis (Hwang & Cox, 2024). To determine the effect of dietary CHX on Clu bliss particle dynamics *in vivo*, we fixed and immunolabeled ovarioles from CHX-fed females (Fig 5D-F”). As expected, untreated well-fed females displayed robust Clu particles (Fig 5D, D”). Like our *ex vivo* experiment, CHX-fed females also exhibited robust Clu particles, indicating that CHX feeding does not disassemble particles (Fig 5E, E”, F, F”). We previously demonstrated that mitochondria mislocalize in nurse cells when the females are exposed to various stressors (Cox & Spradling, 2009; Sen & Cox, 2016; Sen et al., 2015; Sheard et al., 2020). CHX-feeding not only maintained bliss particles, but also maintained mitochondrial morphology and distribution, supporting that feeding CHX was not stressful for the nurse cells (Fig 5E-F’’).

Since CHX did not disassemble bliss particles *ex vivo* and *in vivo*, we wanted to test the effect of CHX on particle assembly. To do this, we dissected ovaries from starved *clu^CA06604^* females that have completely disassembled bliss particles (Fig 5H, I, S8-9 Movie) (Sheard et al., 2020). Normally, adding insulin to the media causes bliss particles to quickly assemble (Fig 5H, H’, S8 Movie, Sheard 2020). However, with preincubation of CHX, insulin-induced bliss particle assembly was blocked (Fig 5G, I, I’, S9 Movie). Taken together, these data suggest that the stalled ribosomes and increased polysomes occurring with CHX treatment do not disassemble already formed bliss particles. However, insulin signaling is insufficient to drive particle assembly when ribosomes are stalled in the absence of particles.

### Cycloheximide maintains bliss particles in the presence of nutritional stress

For effective *Drosophila ex vivo* imaging, insulin must be added to the media to fully support the tissue (Morris & Spradling, 2011). If it is omitted, egg chamber development is not normal, and the samples start to degenerate (Prasad et al., 2007; Prasad & Montell, 2007). Since *Drosophila* Insulin-like peptides secreted by the brain are required for normal egg chamber development, incubating follicles without insulin does not supply effective nutritional signaling and is stressful to the tissue (LaFever & Drummond-Barbosa, 2005; Richard et al., 2005). Since CHX did not abolish bliss particles, in contrast to PUR, we wondered whether CHX treatment could confer a protective effect to maintain bliss particles in the presence of stress. Ovarioles, which were dissected from well-fed *clu^CA06604^* females and pre-incubated in insulin-free media (CS), disassembled bliss particles within 30 minutes, supporting the critical role of nutrition in maintaining particles (Fig 6B, B’). Surprisingly, adding CHX *ex vivo* after dissection was sufficient to maintain bliss particles for 30 minutes, even in the absence of insulin (Fig 6B”). To test whether CHX could also protect bliss particles from nutritional stress *in vivo*, we starved flies after feeding CHX. Well-fed *w^1118^* females were switched to wet yeast paste containing CHX and fed for 24 hours. They were then starved on water only for 5 hours. Five hour starvation completely disassembled bliss particles (Fig 6C, C”) (Sheard et al., 2020). However, providing CHX for 24 hours before starvation was sufficient to maintain bliss particles and protect them from disassembly (Fig 6D, D”, E, E”). Not only are particles maintained, CHX feeding before starvation appears to decrease cellular nutritional stress as indicated by normal mitochondrial localization (Fig 6C’ vs D’, E’) (Sheard et al., 2020). This observation supports that stalled ribosomes and increased polysomes maintain and protect bliss particles even with decreased nutrition *ex vivo* and *in vivo*. Decreased protein synthesis also appears to protect the nurse cells from nutritional stress-induced mitochondrial mislocalization.

**Fig 6.**
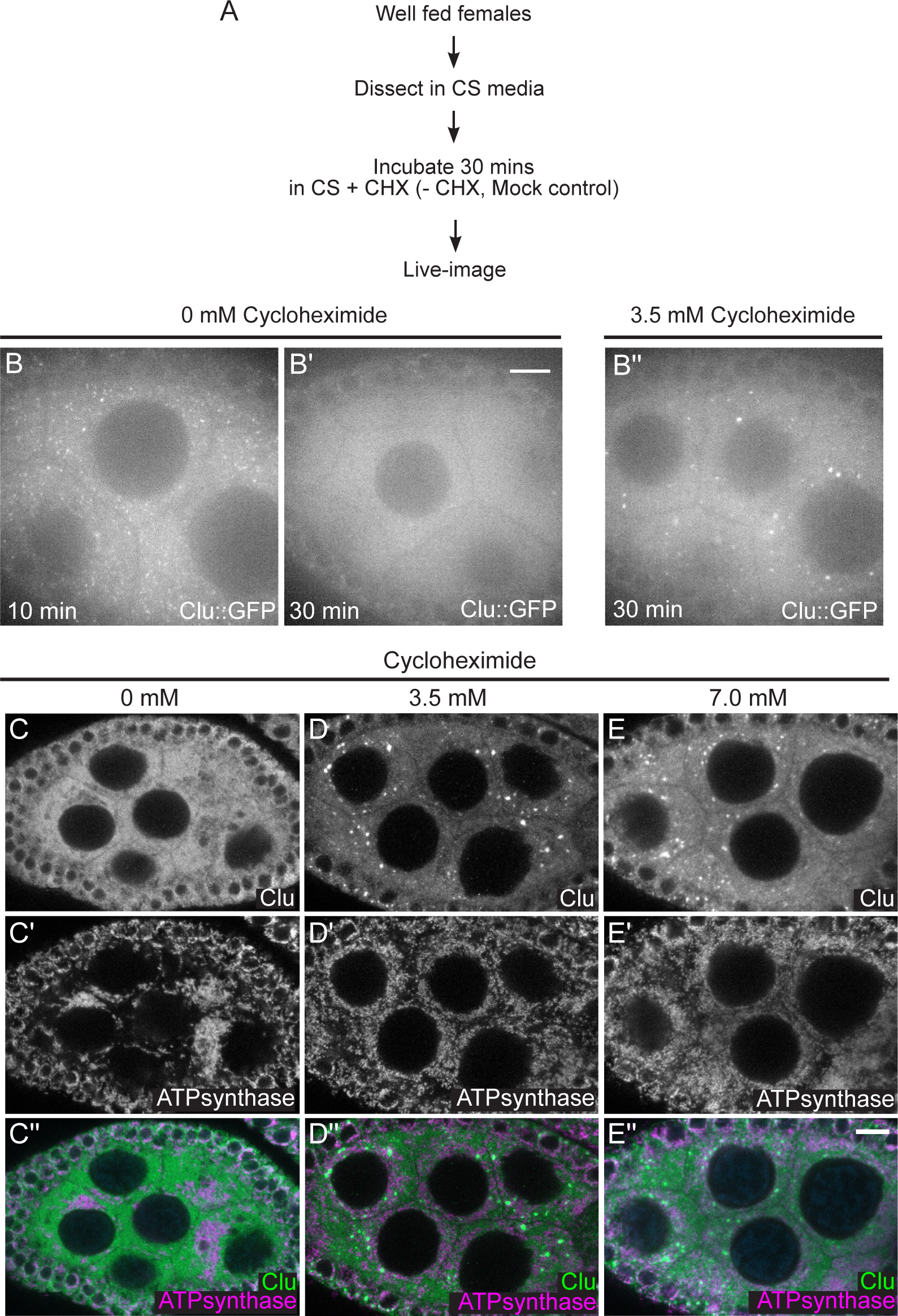
Cycloheximide maintains bliss particles in the presence of nutritional stress. (A) Workflow for the experiment. Ovarioles from well-fed *clu^CA06604^* females were dissected without insulin, then treated with 3.5 mM CHX (B”) or not (mock treatment, B, B’). (B) Still-image of stage 7 follicle from well-fed females maintains Clu particles without insulin in 10 minutes (n= 5 follicles, 85% of the nurse cells having Clu::GFP particles, see S2 Table for details), (B’) but lost Clu particles in 30 minutes after dissection (n= 5 follicles, 42% of nurse cells having Clu::GFP particles, see S2 Table for details). (B”) Still images of stage 7 follicle treated with 3.5 mM CHX for 30 minutes demonstrating CHX treatment does not cause particle dispersion even without insulin (n=10 follicles, 98% of nurse cells having Clu::GFP particles, see S2 Table for details). The focal plane was selected to show at least three to four nuclei were clearly visible in the nurse cells, with approximately 25% depth from the top surface of each follicle (See Materials and Methods for details). The follicle stages analyzed (n): no CHX in 10 mins, stage 5 (3), stage 6 (0), stage 7 (2); no CHX in 30 mins, stage 6 (1), stage7 (3), stage8 (1); 3.5 mM CHX, stage 5 (3). Stage 6 (3), stage 7 (4). More details, including the number of nurse cells having Clu particles and the number of dissected animals, are described in S2 Table. (C-E”) Immunostaining of stage 7 follicles from well-fed *w^1118^* females subsequently starved for 5 hours after feeding for 24 hours with wet yeast paste containing (C-C”) 0 mM, (D-D”) 3.5 mM, and (E-E”) 7 mM. Starvation disrupts particles (C) and causes mitochondrial clustering (C’) as we previously showed (Sheard 2020). (D-D”) 3.5 mM and (E-E”) 7 mM CHX feeding does not disperse Clu bliss particles (D, E) nor cause mitochondrial clump (D’,E’) even with starvation. 87% (3.5 mM CHX following starvation, n=15 follicles), and 94% (7 mM CHX following starvation, n=14 follicles) of nurse cells from the observed follicles had Clu particles (See S2 Table for details). Images are 2 µm projections assembled from 0.42 µm sections. The focal plane was chosen to show at least more than three nurse cells having clear visibility for nulear and cytoplasmic area but also to avoid dim fluorescence signals due to deeper depth (See Materials and Methods for details). The follicle stages examined (n) for each condition: no CHX-5 hour starvation, stage 5 (2), stage 6 (6), stage 7 (3), stage 8 (3); 3.5 mM CHX-5 hour starvation, stage 5 (3), stage 6 (2), stage 7 (6), stage8 (4); 7 mM CHX-5 hour starvation, stage5 (1), stage 6 (3), stage 7 (4), stage 8 (2). More details, including the number of nurse cells having Clu particles and clumped mitochondria and the number of dissected animals, are described in S2 Table. (C, D, E) White = anti-Clu. (C’, D’, E’) White = anti-ATP synthase. (C”, D”, E”, merge) Green = anti-Clu, magenta = anti-ATP synthase. Scale bar = 10 µm in B” for B-B” and in E” for C-E”.

### Cycloheximide maintains bliss particles in the presence of oxidative stress

Hydrogen peroxide (H_2_O_2_) is highly toxic to cells, immediately causing a sharp increase in reactive oxygen species and oxidative damage (Vona et al., 2021). Previously, we demonstrated that adding H_2_O_2_ to cultured ovarioles quickly disassembles Clu bliss particles (Fig 7B-B”’, S10 Movie) (Sheard et al., 2020). Since CHX treatment of cultured ovarioles maintained Clu bliss particles even in the absence of insulin (Fig 6B”), we tested if CHX treatment could maintain bliss particles with high levels of oxidative stress (Fig 7). We cultured ovaries from well-fed *clu^CA06604^* females, added CHX, and then exposed the ovarioles to H_2_O_2_ (Fig 7A). Surprisingly, adding CHX maintained Clu bliss particles in the presence of H_2_O_2_ (Fig 7C-C”’, S11 Movie). This was also observed for a higher CHX concentration (Fig 7D-D”’, S12 Movie). This observation suggests that CHX treatment and increased polysomes are sufficient to maintain and protect Clu bliss particles from oxidative stress-induced particle disassembly, in addition to protection from nutritional stress.

**Fig 7.**
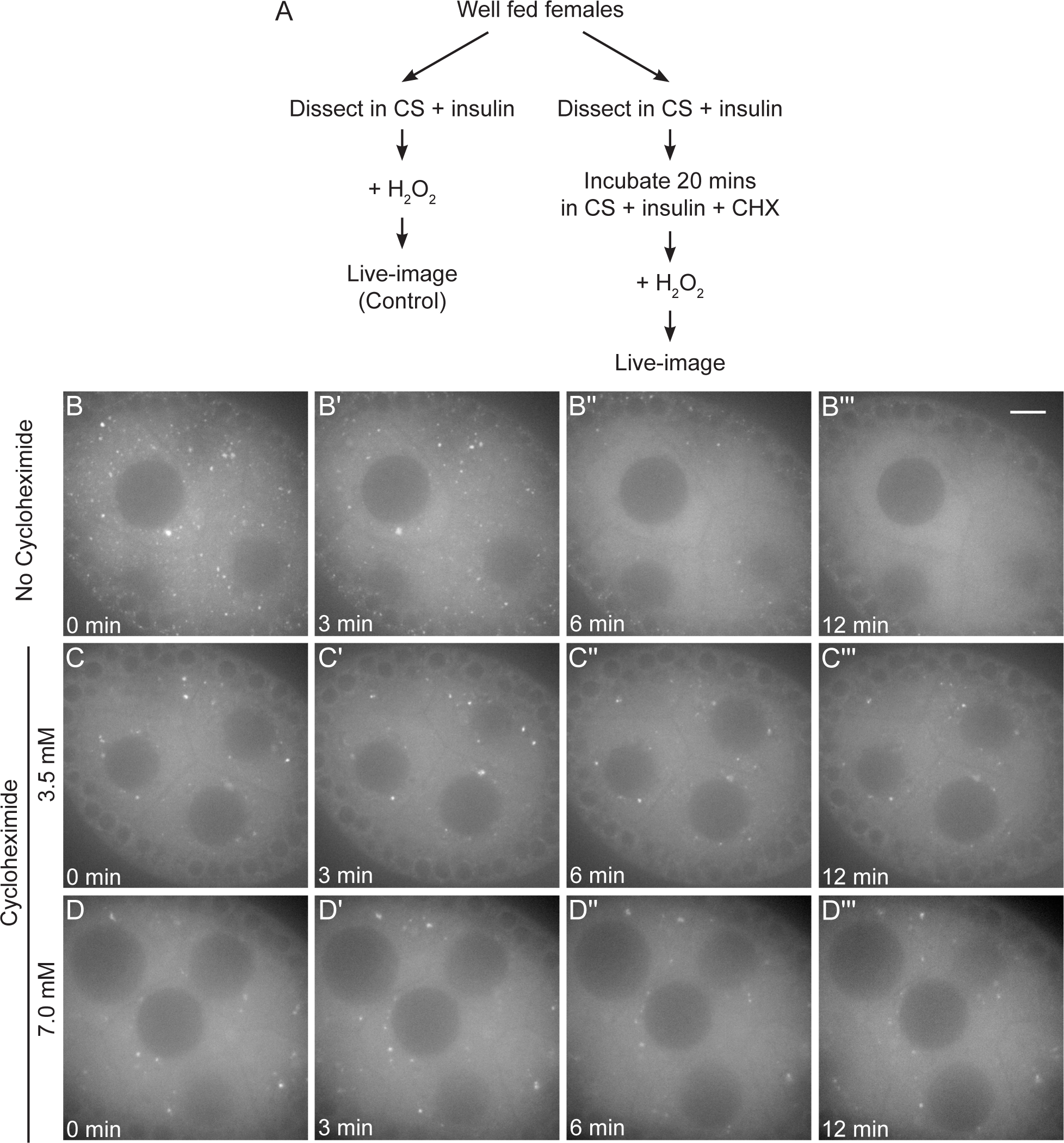
Cycloheximide maintains bliss particles in the presence of oxidative stress. (A) Workflow for the experiment. Ovarioles dissected from well-fed *clu^CA06604^* females were treated with CHX, then exposed to 2 mM hydrogen peroxide (C-D’”). (B-B”’) Still-images of stage 7 follicle from S10 Movie showing the addition of 2 mM hydrogen peroxide disperses bliss particles as we previously showed (Sheard 2020). 93% of nurse cells (n=9 follicles, see S2 Table for details, 2020) clearly showed dispersion of bliss particles after hydrogen peroxide treatment. (C-C”’) Still-image of stage 7 follicle from S11 and (D-D”’) S12 Movies showing CHX treatment protects bliss particles from oxidative stress-induced dispersion. None of the nurse cells pre-treated with 3.5 mM CHX for 20 minutes (n=9/9 follicles) showed dispersion of bliss particles after hydrogen peroxide treatment, and 13% of the nurse cells pre-treated with 7 mM CHX for 20 minutes (n=7 follicles, see S2 Table for details) showed dispersion of CluGFP particles after hydrogen peroxide treatment. The focal plane was chosen to show at least three to four nurse cells having clear visibility of nuclear and cytoplasmic area, with approximately 25 % depth from the top surface of each follicle (See Materials and Methods for details). Follicle stages analyzed (n): no CHX-hydrogen peroxide, stage 5 (3), stage 6 (2), stage 7 (3), stage 8 (1); 3.5 mM CHX-hydrogen peroxide, stage 5 (1) stage 6 (1), stage 7 (5), stage 8 (2); 7 mM CHX-hydrogen peroxide, stage 5 (2), stage6 (3), stage7 (2). S2 Table. Scale bar = 10 µm in B”’ for B-D”’.

### Clu particles associated with mitochondria move more slowly

Because we observed that CHX treatment had an impact on bliss particle assembly and disassembly, we tested whether CHX treatment affects bliss particle velocity (Fig 8, S6-7 Movies). Using kymographic analysis, we tracked the movement of the directed Clu particles and measured their velocities (Fig 8B, B’, white arrows). The average particle velocities were not affected by CHX treatment (Fig 8C), indicating that modulating translation activity by CHX did not impede particle movement although CHX affected bliss particle assembly/disassembly. Previously, we demonstrated that Clu particles are juxtaposed to, but not overlapping with, mitochondria in fixed tissues (Cox & Spradling, 2009). However, we did not investigate whether this mitochondrial association is related to particle movement. To answer this question, we live-imaged follicles dissected from *clu^CA06604^* females and treated them with the mitochondrial membrane potential sensitive dye tetramethylrhodamine ethyl ester (TMRE) to visualize mitochondria. Clu particles frequently moved in the cytoplasm while associated with mitochondria (Fig 9A-E, S13 Movie). Using kymographic analysis, we tracked the speed of mitochondrial-associated and unassociated Clu particles (Fig 9B, C, E) and measured particle size (Fig 9A, D, E). The Clu particles associated with mitochondria moved significantly slower than the particles not associated (Fig 9C). Particle size did not correlate with whether or not the particle was associated with a mitochondrion (Fig 9D). However, as particle size increased, the speed decreased (Pearson correlation, *r* = -0.3287, Fig 9E). To confirm that particle size did not affect mitochondrial association, we performed logistic regression analysis (Fig 9F, G, S2B, C). Using simple logistic regression analysis (one independent variable: size or speed), we found that the speed of particle movement greatly affected mitochondrial association, but Clu particle size did not affect whether the particle was mitochondria-associated (Fig 9F vs. G, S2B vs. C). This suggests that the speed of the particle, not the size, determines whether there is particle-mitochondrion association.

**Fig 8.**
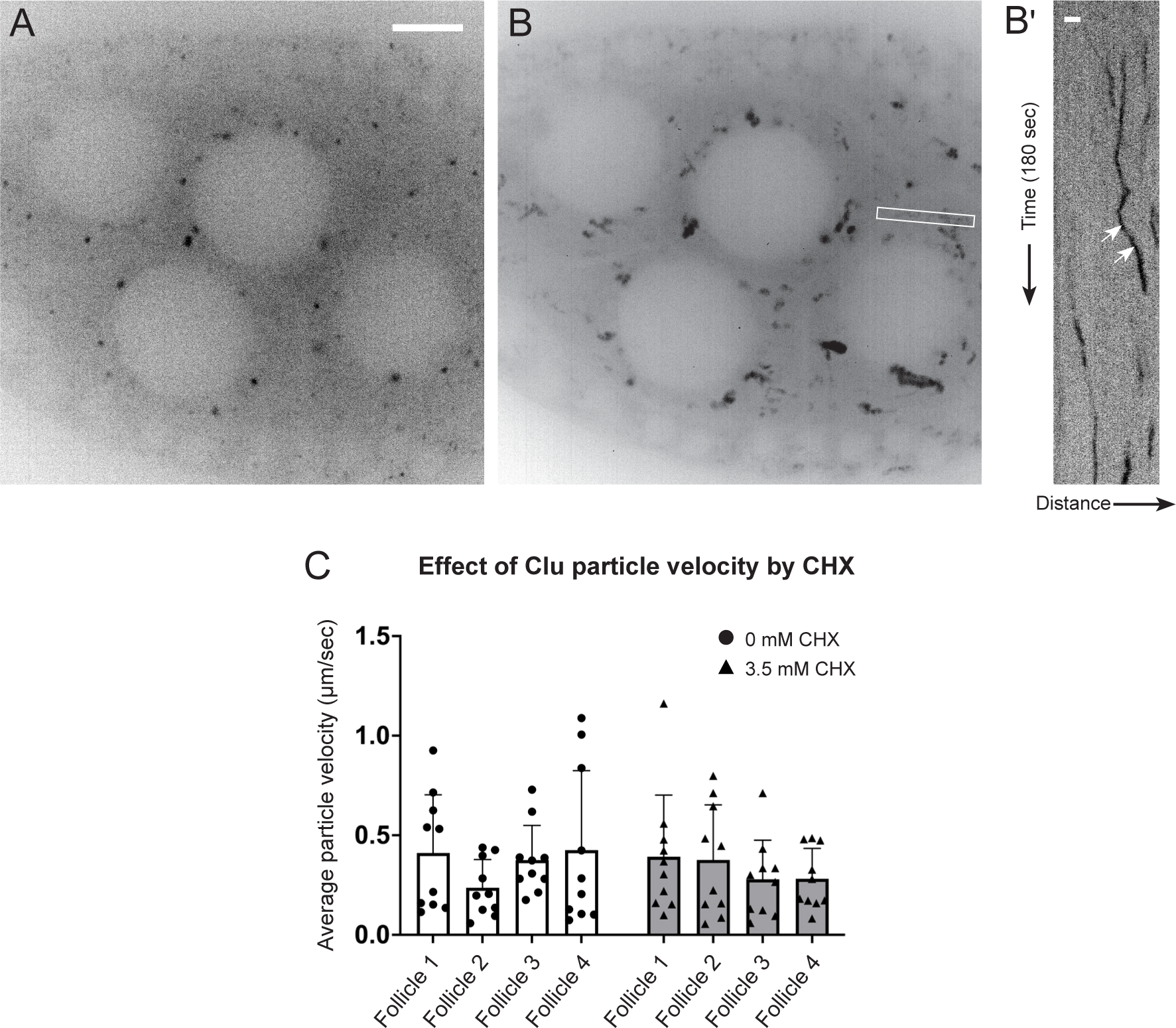
Cycloheximide does not affect velocity of Clu particle. (A) Representative still-image of Clu::GFP particles from S6 Movie. This still-image is Fig 5C, inverted in LUT. (B) Three-minute S6 movie was stacked to find the orientations of Clu particles. The thin white box shows the area used to make a kymograph in (B’). (B’) Representative kymographs of Clu particles. The white arrows indicate a directed movement of Clu particles and the velocity was measured by applying a straight line using ImageJ. More details for velocity calculation are described in S2 Table. (C) No effect of cycloheximide on the average velocity of bliss particles. Dots in the graph represent the velocity of each particle. Six particles were analyzed in each follicle. The graph was generated using GraphPad Prism. Follicle stages analyzed (n): no CHX (control), stage 6 (1), stage 7(2), stage 8 (1); 3.5 mM CHX, stage 6 (1), stage 7 (3). Scale bar = 10 µm in A for A and B. Scale bar 2 µm in B’.

**Fig 9.**
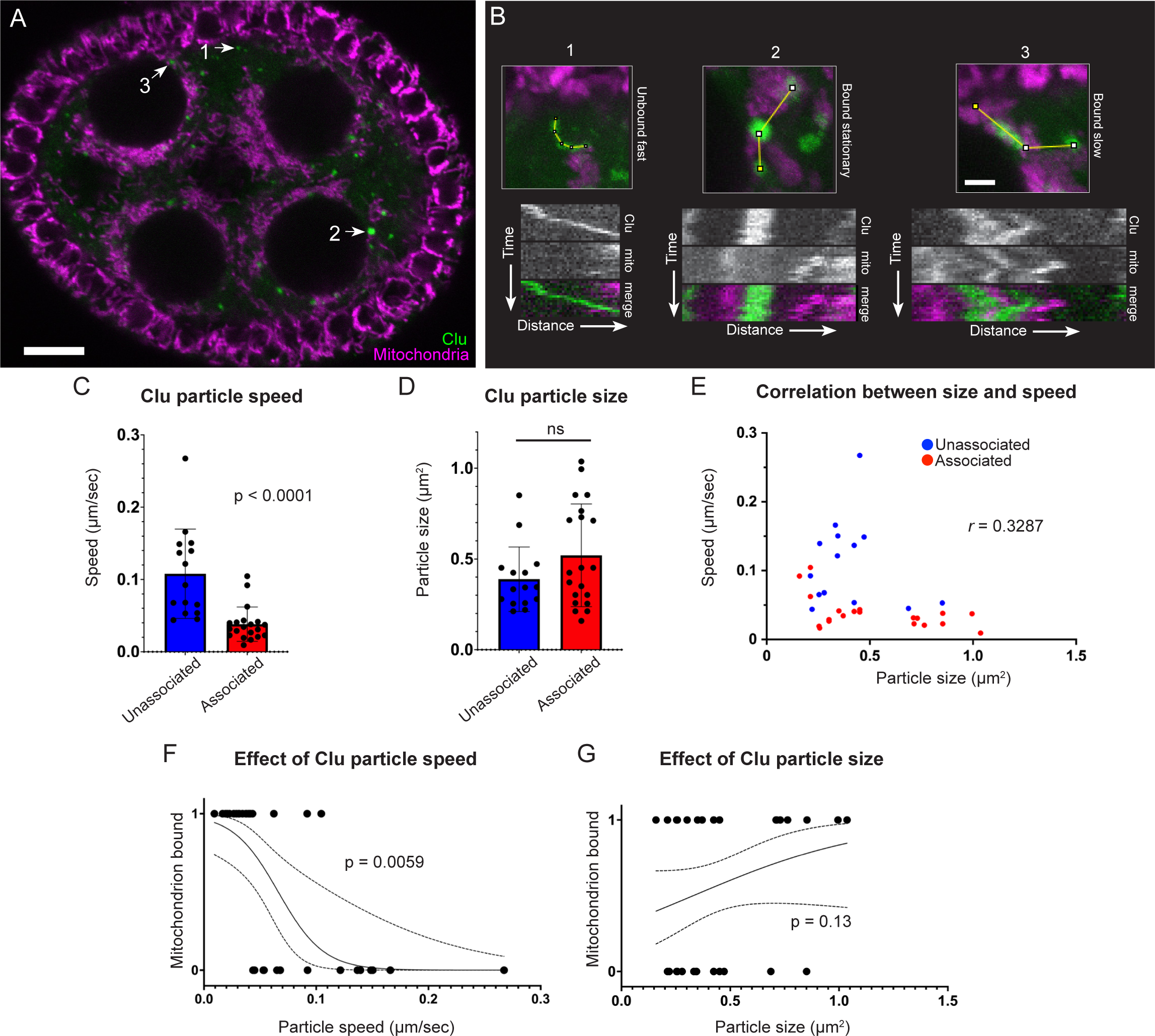
Clu particle associated with mitochondria moves slowly. (A) Still-image (S13 Movie) of a follicle from a Clu GFPTrap *clu^CA06604^* female labeled with 50 nM TMRE. (B) Representative kymographs of Clu particles (A, arrows). Kymographs (lower panels) were obtained from the z-stacked image of frames 70-85 (for 1 minute) of S13 Movie by applying segmented lines (upper panels). Representative graph of (C) speed and (D) size of Clu particles with or without mitochondrial association. A bar graph represents an arithmetic mean. Unpaired t-test was performed for statistical significance. (E) Representative graph of Clu particles plotted by speed, size, and mitochondrial association. Pearson correlation coefficient was obtained to measure the relationship between size and speed. (F, G) Simple logistic regression analysis of (E). This predicts a probability of mitochondrial association depending on particle speed (F) or size (G). The value 1 of the y-axis represents the mitochondrially-associated particle, and 0 represents the particles free from mitochondria. The solid line indicates the mean of the probability, and the dotted line indicates a 95% confidence interval. Logistic regression tests were performed using GraphPad Prism. The total number of particles (n) = 35. The number of follicles examined is as follows: stage 5 (2), stage 6 (4), and stage 7(3). More details, including the number of particles in each follicle, particle area, particle speed, and particle binding are described in S3 Table. Red = mitochondria-associated particles, blue = mitochondria-unassociated particles. (A, B) Green = Clu, magenta = TMRE. Scale bar = 10 µm in A. Scale bar = 2 µm in B for upper panels.

## Discussion

### Clu bliss particle assembly requires regulated levels of functional Clu

Clu could act as a scaffold protein within particles as this is a common feature for cytoplasmic granule proteins (Buchan, 2014). Many cytoplasmic bodies and granules have associated proteins containing low-complexity or prion-like domains. Clu’s large “M” domain that is predicted to be unstructured but does not contain canonical intrinsically disordered motifs. It could still be important for particle dynamics, but this has yet to be tested. We previously demonstrated that Clu can self-associate, but we do not yet know whether Clu forms dimers, multimers, or self-association is due to protein aggregation in bliss particles (Sen & Cox, 2016). Determining which domains are required for self-association using ectopic Clu expression has proved challenging *in vivo* since endogenous Clu must be absent and *clu* null mutants are quite sick. We have identified that the DUF, Clu, and TPR domains are required for particle assembly and for association with endogenous bliss particles but we do not yet know whether they play a role in molecular self-association in *Drosophila*. The TPR domain of CLUH has been reported to be necessary for CLUH self-association, although it appears that the self-association is not direct, suggesting CLUH forms multimers (Hemono et al., 2022). In addition, we previously showed that, in *Drosophila*, the TPR domain is essential for mRNA association (Sen & Cox, 2016), which was confirmed to be the case for CLUH (Hemono et al., 2022). Ectopically expressing full length (FL)-Clu at high levels caused bliss particles to disassemble, which differs from the observation of CLUH granules (discussed below (Pla-Martin et al., 2020)). While this could be due to toxicity, it seems unlikely since overexpression of ΔTPR and ΔDUF did not cause endogenous particle disassembly, nor did overexpression of an unrelated protein CPB. An alternative explanation could be that high concentrations of FL-Clu disrupt required stoichiometry for forming bliss particles. Too much functional Clu could disrupt potential liquid-liquid phase separation that may regulate bliss particle assembly and disassembly, or the increased protein could exert a dominant effect by sequestering factors necessary for particle assembly.

### Assembly and maintenance of Clu bliss particles require polysomes

As Clu associates with mRNA and ribosomal proteins, we tested the effect of translation inhibitors on assembly and disassembly with and without stress. CHX and PUR treatment are powerful tools to assess how stalled translation affects the dynamics of particles and granules involved in the posttranscriptional regulation of mRNA. PUR decreases the amount of polysomes, terminating translation, releasing ribosomes, and increasing the amount of messenger RNPs (mRNPs). In contrast, CHX prevents elongation. This results in an inhibition of polysome to mRNP conversion and thus increases stalled ribosomes and decreases levels of mRNPs. Stress granules are composed of mRNPs, stalled preinitiation complexes, and other proteins involved in translation. Under normal conditions, low concentrations of puromycin rarely assemble stress granules, but longer incubation with higher concentrations does (Bounedjah et al., 2014; Ihn et al., 2024; Kedersha et al., 2000; Martinez et al., 2016). In the presence of stress, puromycin increases the number of stress granules, whereas cycloheximide disassembles them. This dynamic is caused by an increased number of mRNPs available with puromycin and a decreased number available with cycloheximide. P-bodies generally contain proteins that are associated with mRNA decay or silencing. P-body maintenance depends on the presence of translationally repressed mRNA. In the presence of CHX, mRNA trapped in polysomes causes P-body loss. However, with PUR treatment, the number and size of P-bodies increase due to the increase in non-translatable mRNPs (Eulalio et al., 2007; Patel et al., 2016).

By treating follicles with PUR and CHX, we found that the availability of polysomes regulates assembly and disassembly of bliss particles. Administering PUR *ex vivo* rapidly disassembled bliss particles, whereas CHX had no effect *ex vivo* and *in vivo*. This suggests that increasing the amount of mRNPs and decreasing polysomes disassembles bliss particles. When particles are absent, CHX treatment blocked insulin-induced assembly *ex vivo* suggesting that increased polysomes alone are insufficient for Clu particle assembly. The increased polysomes might be physically prevented from separating into Clu particles despite the unchanged Clu levels. This also suggests that insulin signaling, which initiates a strong signaling cascade, cannot overcome the CHX block in assembly. Nutritional and oxidative stress disassemble Clu bliss particles. Both stressors cause decreased translation rates through integrated stress response signaling. During nutritional stress *ex vivo* and *in vivo*, CHX treatment maintained Clu particle assembly, suggesting that, once particles are formed, polysomes present in Clu particles stabilize the particles. This was also true for oxidative damage *ex vivo* caused by hydrogen peroxide. Together, these data support a model whereby *Drosophila* Clu bliss particles harbor actively translating mRNAs and particle assembly relies on the presence of polysomes. Evidence for Clu bliss particles as sites of active translation is supported by observations of CLUH granules (Pla-Martin et al., 2020).

### Comparing *Drosophila* Clu and vertebrate CLUH subcellular localization

Clu forms large, prominent, and highly dynamic particles in germ cell cytoplasm (Cox & Spradling, 2009; Sheard et al., 2020). These particles are found in *Drosophila* somatic tissues as well (Sen et al., 2013; Sheard et al., 2020; Wang et al., 2015). We previously demonstrated that Clu particles are highly sensitive to stress. Using live-imaging, Clu bliss particles quickly disassemble in the presence of hydrogen peroxide, and for germ cells lacking particles due to starvation, adding insulin to the media causes particle assembly in minutes (Sheard et al., 2020). Nutritional stress *in vivo* from starvation causes particle disassembly, with no decrease in protein, and particles reassemble upon feeding (Sheard et al., 2020). Various studies in vertebrate systems have demonstrated CLUH localization in the cell. In COS7 cells, CLUH exhibits a granular pattern, particularly after Triton-X 100 extraction (Gao et al., 2014). In primary hepatocytes and HeLa cells, CLUH is reported to form granules that colocalize with some, but not all, components of stress granules, and these granules increase with stress (Pla-Martin et al., 2020). In addition, in contrast to the data shown here, CLUH overexpression induced the formation of peripheral CLUH granules in about 40% of transfected HeLa cells (Pla-Martin et al., 2020). However, additional reports have shown CLUH is broadly diffuse in the cytoplasm in HCT116 cells and CLUH does not localize with the stress granule component G3BP1 (Hemono et al., 2022). In agreement with our observation, CLUH granules did not disassemble in response to CHX treatment, although these granules were the peripheral granules assembled by overexpression of CLUH (Pla-Martin et al., 2020). Finally, two groups have shown CLUH colocalizes with one or two structures composed of SPAG5/Astrin, which is a mitotic spindle protein during mitosis that localizes to microtubule plus-ends in the cytoplasm during interphase (Dunsch et al., 2011; Hemono et al., 2022; Schatton et al., 2022). It is not clear at present why insects’ Clu localization and dynamics are so different from vertebrate CLUH. One possibility could be that vertebrates may simply have different CLUH dynamics from Drosophila Clu due to differences in cell types and species. Another possibility could be that Clu particles may act differently *in vivo* compared to CLUH in cell culture experiments due to differences in cell physiology.

### Nurse cells with CHX-stabilized Clu particles have reduced cellular stress as indicated by mitochondrial localization

Starvation causes germ cell mitochondria to cluster (Sheard et al., 2020). This occurs not only with nutritional stress and appears to be a hallmark of cellular stress. When bliss particles are absent, mitochondria are clumped, but when the particles are present, mitochondria disperse throughout the germ cell cytoplasm in the normal pattern. This indicates that normal mitochondrial localization highly correlates with the presence of bliss particles (Cox & Spradling, 2009; Sen et al., 2015; Sheard et al., 2020). We found similar mitochondrial dynamics with CHX treatment. Females fed a rich CHX diet maintained assembled bliss particles and mitochondrial distribution (Fig 5D-F”). Bliss particles that remained assembled with CHX treatment followed by starvation also had normal mitochondrial localization (Fig 6C-E”). This could be attributed to CHX increasing the level of amino acids and thus stimulating TOR activity, a downstream component of the insulin signaling pathway (Beugnet et al., 2003). In addition, Clu particle association with mitochondria decreased particle speed. However, we are unable to distinguish whether the relationship between particle speed and mitochondrial association is because particles physically slow due to the complex size, or whether mitochondria can only associate with slower particles. We were unable to determine whether particle-mitochondrion association affected mitochondrial activity at the single organelle level using TMRE due to variability and efficiency of staining.

In this study, we identified three protein domains of Clu that are necessary to assemble Clu bliss particles. We also found that overexpression of functional FL-Clu causes particle disassembly. In addition, we found that Clu bliss particles require the presence of polysomes to remain assembled and that stabilizing polysomes protects bliss particles from disassembly. Since Clu associates with ribosomes and is a ribonucleoprotein, this supports a model whereby Clu bliss particles are active sites of translation under non-stressed conditions but disassemble when cellular stress increases. Since Clu is closely tied to mitochondrial function and binds mRNAs encoding mitochondrial proteins, particle dynamics likely affects mitochondrial function. Disassembly of particles could lead to decreased translation of the associated mRNAs, fewer mitochondrial proteins actively translated in the cytoplasm, and a shift in metabolism in response to stress. There is evidence supporting this idea through studies on CLUH (Pla-Martin et al., 2020). For a better understanding of *Drosophila* bliss particles, several challenges must be overcome. There are many proteins known to associate with P-bodies and stress granules (Anderson & Kedersha, 2006; Ivanov et al., 2019). While we and others have identified Clu/CLUH-associated proteins using coimmunoprecipitation and mass spectrometry, we have yet to localize any of them to bliss particles. In addition, we have yet to determine whether bliss particles are active sites of translation. Particles are easily seen in the nurse cells, yet using fluorescent in situ hybridization is challenging. Nonetheless, given Clu’s critical role in mitochondrial function, fully understanding how these unique and novel RNP particles function will ultimately deepen our knowledge of how mitochondria respond to stress.

## Materials and Methods

### Fly stocks

Fly stocks were maintained on standard cornmeal fly media. Animals were grown at room temperature. The following stocks were used for experiments: *w^1118^*, *w^1118^; clu^CA06604^* (Cox & Spradling, 2009), *w\**; *clu^CA06604^*/*CyO; nanosGAL4*/*TM3 Sb*, *w^1118^*; *UASp-cluΔDUF*::*mScarlet*/*TM3 Sb*, *w^1118^*; *UASp-cluΔClu*::*mScarlet*/*TM3 Sb*, *w^1118^*; UASp-cluΔTPR::*mScarlet*/*TM3 Sb*, *w^1118^*; *UASp-FLclu*::*mScarlet*/*TM3 Sb*, *w\**; *Kr^If-1^*/*CyO*; *P{w[+mW.hs]=GAL4-da.G32}UH1* (Bloomington Drosophila Stock Center (BDSC), Bloomington, IN, USA, BSC# 55850), and *w\**; *M{w[+mC]=UASp-mCherry.cpb}ZH-86Fb*/*TM3 Sb^1^* (BDSC, Bloomington, IN, USA, BSC# 58728). A newly eclosed fly is considered day 0.

### Transgenic flies and constructs

For the C-terminal fusion of mScarlet for live-imaging, we created a Gateway destination vector pPgateWmScarlet-i for subcloning. Briefly, the mScarlet-i coding sequence was amplified from pCytERM_mScarlet-i_N1 (Addgene, Watertown, MA, USA, cat#. 85068) using the following primers, 5’-TAG GCC ACT AGT GTG AGC AAG GGC GAG GCA GT-3’and 5’-TGC TTA GGA TCC TTA CTT GTA CAG CTC GTC CA-3’. The amplicons were subcloned into a pQUASp (Addgene, Watertown, MA, USA, cat#. 46162), placing the UASp promoter upstream of the mScarlet-i coding sequence. Ampicillin resistance was used to select positive clones, which were verified by restriction digest and sequencing. The resulting pUASp-mScarlet-i construct was converted to a Gateway destination vector using the Gateway™ Vector Conversion System (Invitrogen, Waltham, MA USA, cat#. 11828029). Chloramphenicol resistance was used to select positive clones, which were verified by sequencing. The pQUASp vector was a gift from Christopher Potter (Addgene plasmid #46162; http://n2tnet/addgene:46162; RRID: Addgene_46162), and pCytERM_mScarlet-i_N1 vector was a gift from Dorus Gadella (Addgene plasmid # 85068; http://n2t.net/addgene:85068; RRID: Addgene_85068) (Bindels et al., 2017). Gateway entry vectors with full-length Clu (Clu_pENTR) or domain-deleted Clu (ΔDUF_pENTR, ΔClu_pENTR, and ΔTPR_pENTR) were previously described (Sen & Cox, 2016). Each entry vector was cloned into pPgateWmScarlet-i using LR Clonase mix (Invitrogen, Waltham, MA, USA, cat#. 11791020) following the manufacturer’s directions. The resulting expression vectors were selected by ampicillin resistance and verified by sequencing. For transgenic flies, the vectors were commercially injected (BestGene Inc. Chino Hills, CA, USA).

### Live-imaging for analysis of Clu particle dynamics

Live-imaging with ovarioles was performed as previously described (Sheard et al., 2020) with some modifications. Ovaries were dissected in a live-imaging media composed of Complete Schneider’s (CS) media and 200 µg/mL of insulin. The CS media was Schneider’s Drosophila medium (Fisher Scientific, Hampton, NH, USA, cat#. BW04351Q) supplemented with 15% heat-inactivated fetal bovine serum (CPS Serum, Parkville, MO, USA, cat#. FBS-500HI) and penicillin (100 U/mL) - streptomycin (100 µg/mL) (Fisher Scientific, Hampton, NH, USA, cat#. BW17602E). After separating ovarioles from an ovary and eliminating the connected muscle sheath and nerve fibers, the tissues were transferred into a 35 mm MatTek glass bottom dish (MatTek Corporation, Ashland, MA, USA, cat#. P35G-0-20-C) with a live-imaging media. To simultaneously visualize Clu particles and mitochondria, TMRE (tetramethylrhodamine, ethyl ester, perchlorate) (Anaspec Inc, Fremont, CA, USA, cat#. AS88061) was diluted to 50 nM in the dissection media. Tissues were incubated for 20 minutes and imaged without washing. The follicles were mainly chosen between stages 5 to 7, previtellogenic stages, for better observation with nurse cells. The focal plane was selected to have at least three to four nurse cells with a clear visibility of nuclear and cytoplasmic areas, with appproximately 25% depth from the top surface of a follicle. Follicle stages were determined with a size of a follicle as described in the reference (Spradling, 1993). More details for each experiment, including the number of dissected animals and replicates are describe in supplementary tables. Live images were obtained using a Nikon A1 plus Piezo Z Drive Confocal microscope at 60x (Nikon Corporation, Tokyo, Japan) or a Nikon Eclipse Ti2 spinning disk microscope at 100x (Nikon Corporation, Tokyo, Japan). Fiji ImageJ was utilized to analyze confocal images (Schindelin et al., 2012).

### Live-imaging for Clu particle dynamics with puromycin, cycloheximide and hydrogen peroxide treatment

The working solution for each chemical was prepared just before performing an experiment as follows: 10 mg/mL puromycin (Gibco^TM^, Waltham, MA, USA, cat#. A1113803) was diluted to 20 µM in a media composed of Complete Schneider’s (CS) media and 100 µg/mL of insulin (CS/Ins); cycloheximide powder (CHX, Sigma-Aldrich, Burlington, MA, USA, cat#. C7698) was dissolved in CS for 14 mM stock solution, which was further diluted to 3.5 mM and 7 mM CHX-containing CS or CS/Ins media; and 30% hydrogen peroxide (Sigma-Aldrich, Burlington, MA, USA, cat#. H1009) was diluted to 4% hydrogen peroxide in CS/Ins or CHX-containing CS/Ins. Ovaries were dissected as described above with corresponding media, CS or CS/Ins, depending on the purpose of each experiment. To test the puromycin effect on bliss particle dynamics, dissected ovaries with CS/Ins media were transferred into a 35 mm MatTek glass bottom dish with 50 µL of CS/Ins media, and live imaging was performed after adding 50 µL of CS/Ins containing 20 µM puromycin to the ovaries to make a final concentration of 10 µM puromycin. To test the CHX effect on bliss particle dispersion, dissected ovaries with CS/Ins media were incubated for 20 minutes with 3.5 mM CHX-containing CS/Ins (CS/Ins/CHX) following twice wash with the same media, and then a live-imaging was performed. To test the CHX effect on bliss particle formation, ovaries were dissected from starved animals in CS, incubated for 20 minutes with 3.5 mM CHX-containing CS (CS/CHX) following twice wash with the same media, re-washed twice with CS/CHX containing 100 µg/mL insulin (CS/CHX/Ins), and then switched to CS/CHX/Ins for immediate live-imaging. To test the CHX effect on particle dispersion without insulin, ovaries were dissected from *w^1118^*; *clu^CA06604^* in CS, washed twice with CS containing 3.5 mM CHX (CS/CHX), incubated for 30 minutes with the same media, and then live images were obtained. To produce oxidative stress, dissected ovaries were incubated with 50 µL of CS/Ins or CS/CHX/Ins for 20 minutes following twice wash with a corresponding media. Live-imaging was performed after adding 50 µL of each corresponding media containing 4 mM hydrogen peroxide to make a final concentration of 2 mM hydrogen peroxide. Images were obtained using a Nikon A1plus Piezo Z Drive Confocal microscope at 60x or a Nikon Eclipse Ti2 spinning disk microscope at 100x. Selection of follicle stages and a focal plane was performed as described above. More details of each experiment including the number of dissected animals and replicates are described in S2 Table.

### Generating and analysis of kymographs

The still frames of the time-lapse image were stacked to track particle movements. Fiji ImageJ was utilized to generate kymograph and measure velocity with a tool, Multi Kymograph, or Plugins, KymographBuilder (Mary et al., 2016; Schindelin et al., 2012). To measure the velocity of directed Clu particles for CHX treatment, live time-laps images were recorded for 3 minutes and analyzed as follows. Six areas within a follicle were chosen randomly, as shown in Fig 8B, applied with a segmented line to generate a kymograph. A clu particle showing a progressive movement within a kymograph was chosen, and velocity was calculated (more details in S2 Table). To measure the speed of Clu particles of various sizes in relation to mitochondrial association, live time-laps images were recorded for more than 3 minutes after TMRE staining as described above. Clu particle that could be tracked for 1 minute was selected, and the movement of each particle was traced with a segmented line to generate kymograph, which was used to calculate speed (more details in S3 Table). Clu particle-mitochondrial binding was judged by juxtaposing the representative color of Clu (green) and mitochondria (magenta). The size of each Clu particle was calculated using ROI as the area of each particle. Using GraphPad Prism, the graphs were generated and the data was analyzed statistically with unpaired t-test, calculation of Pearson correlation coefficient, and logistic regression (GraphPad Software, Boston, Massachusetts USA, www.graphpad.com).

### Preparation of cycloheximide solution for wet yeast paste to feed flies

CHX was dissolved in water to prepare a 10 mM stock solution that was aliquoted and stored at -20 °C until use. The desired concentration of CHX was prepared by serial dilution of the stock solution. 0.3 g of active dry yeast powder (Red Star® Yeast) was mixed with 450 µL of CHX solution to create a yeast paste which was provided daily as needed.

### Cycloheximide feeding for ovary analysis

Ten female day 0 adult flies and ten male day 0 adult flies were collected in a standard food vial with wet yeast paste, and on day 3, female adults were separated from males. The food vial with fresh wet yeast paste was switched every day to make flies fatten until day 4. On day 4, all female flies in each vial were transferred to a standard food vial containing freshly made yeast paste with CHX. In 24 hours after CHX feeding, fly ovaries were dissected in Grace’s insect media. To determine the effect of starvation after CHX feeding, after feeding CHX for 24 hours, the flies were transferred to an empty vial with a wet piece of tissue with water and maintained for 5 hours. The flies were then dissected and immunostained. The numbers of dissected animals and replicates are described in S2 Table.

### Immunostaining

Ovaries were dissected with Grace’s Insect Medium (Invitrogen, Waltham, MA, USA, cat#11595030) and fixed for 20 minutes (4% paraformaldehyde and 20 mM formic acid in Grace’s Insect Medium). After washing with Antibody wash solution (AWS, 0.1% Triton X-100 and 1% BSA in phosphate-buffered saline) three times for twenty minutes, the tissue was stained with primary antibody overnight at 4 °C. After washing with AWS three times for twenty minutes, the tissues were stained with secondary antibodies overnight at 4 °C, then washed with AWS twice for twenty minutes and stained with 5 ng/mL 4’,6-Diamidino-2-phenylindole (DAPI) solution for ten minutes. After removing the DAPI solution, the tissues were mounted in Vectashield Antifade Mounting Medium (Vector Laboratories, Newark, CA, USA, cat#. H-1000). Images were obtained using a Zeiss LSM 980 confocal laser scanning microscope (Carl Zeiss Microscopy LLC, White Plains, NY, USA). The follicles were mainly chosen between stages 5 to 7, previtellogenic stages, for better observation in nurse cells. The focal plane was selected for ensuring at least three to four nuclei were clearly visible in a nurse cells, with approximately 25% depth from the top surface of a follicle but also to avoid a dim fluorescence signal due to a deeper depth for a fixed imaging. The numbers of dissected animals and replicates are described in supplementary tables. The following antibodies were used: guinea pig anti-Clu N-terminus (1:2000 (Cox & Spradling, 2009)), rat anti-mScarlet-i sdAb-FluoTag-X2 (1:1000, Synaptic System, Goettingen, Germany, cat#. N1302-At488-L), chicken anti-mCherry (1:1000, Novus Biologicals, Centennial, CO, USA, cat#. NBP2-25158), mouse anti-ATP5A1 (1:1000, Abcam, Cambridge, UK, cat#. 14748, or Invitrogen, Waltham, MA, USA, cat#. 439800), goat anti-guinea pig Alexa 488 (1:1000, Invitrogen, Waltham, MA, USA, cat#. A11073), goat anti-guinea pig Alexa 633 (1:1000, Invitrogen, Waltham, MA, USA, cat#. A21105), goat anti-chicken Alexa 568 (1:500, Invitrogen, Waltham, MA, USA, cat#. A11041), goat anti-mouse IgG2b Alexa 488 (1: 500, Invitrogen, Waltham, MA, USA, cat#. A21141), goat anti-mouse IgG2b Alexa 568 (1: 500, Invitrogen, Waltham, MA, USA, cat#. A21144).

## Data availability

All data are contained within the manuscript.

## Supporting information

This Article contains Supporting information

## Acknowledgements

Antibodies obtained from The Developmental Studies Hybridoma Bank, created by the NICHD of the NIH and maintained at The University of Iowa, Department of Biology were used in this study. Reagents were obtained from the Drosophila Genomic Resource Center, Indiana University, Bloomington, IN, supported by NIH grant 2P40OD010949. Stocks obtained from the Bloomington Drosophila Stock Center (NIH P40OD018537) were used in this study.

## Funding and additional information

This work was supported by the National Institutes of Health R01GM127938 to R. T. C. The content is solely the responsibility of the authors and does not necessarily represent the official views of the National Institutes of Health, the Department of Defense, or Uniformed Services University.

## Supporting information

### Supplementary methods

#### Fly stocks

Fly stocks were maintained on standard cornmeal fly media. Animals were grown at room temperature. *w\**; *P{PTT-GA} tral^CA06517^* (Buszczak et al., 2007) were grown at room temperature. A newly eclosed fly is considered day 0.

#### Live-imaging for P-bodies

To test the CHX effect on disaggregation of P-bodies, ovaries were dissected from *w\**; *P{PTT-GA} tral^CA06517^*in CS, and then incubated for 30 minutes with CS containing 3.5 mM CHX (CS/CHX) following twice wash with the same media. Live images were obtained using a Nikon Eclipse Ti2 spinning disk microscope at 100x (Nikon Corporation, Tokyo, Japan). The follicles were mainly chosen in stages 7-8 that show more numbers of P-bodies than earlier stages. The focal plane was selected to have at least three to four nurse cells with a clear visibility of nuclear and cytoplasmic area, with approximately 25% depth from the top surface of a follicle. Changes in P-bodies were determined by a subjective measurement. The numbers of dissected animals and replicates are described in S2 Table.

### Supplementary figure legends

**S1 Fig. Cycloheximide causes reduced sizes and numbers of P-bodies *ex vivo*.** (A) Workflow for the experiment. Well-fed *tral^CA06517^* females were treated with 3.5 mM CHX for 30 minutes, and then live-imaged (B-C’). (B-B’) Still-image of stage 8 follicle showing a mock-control increases P-bodies. 68% of nurse cells had increased numbers and sizes of P-bodies in 30 minutes after mock treatment, and 32% of nurse cells had no changes (n=19 follicles, see S2 Table for details). (C-C’) Still-image of stage 8 follicle showing 3.5 mM CHX treatment decreases P-bodies. 48% of nurse cells had decreased numbers and sizes of P-bodies in 30 minutes after 3.5 mM CHX treatment, 43% of nurse cells had no changes, and 9 % of nurse cells had increased (n=23 follicles, see S2 Table for details). Images are 2 µm projections assembled from 0.5 µm sections. The focal plane was selected to have at least three to four nurse cells with a clear visibility of nuclear and cytoplasmic area, aiming for approximately 25% depth from the top surface of a follicle. Changes in P-bodies were determined by a subjective measurement. Follicle stages analyzed (n): mock control, stage 6 (7), stage 7 (10), stage 8 (2); 3.5 mM CHX, stage 5 (4), stage 6 (5), stage 7 (6), stage 8 (8). More details, including the number of follicles showing changes in the numbers/sizes of P-bodies by CHX treatment and the number of dissected animals, are described in S2 Table. Scale bar = 20 µm in B’ for B-C’.

**S2 Fig: Clu particle dynamics and mitochondria** (A-C) Replicates of the experiment of Fig 9, analysis of Clu particle speed and mitochondrial association. (A) The graph plotted by speed, size, and mitochondrial binding of each Clu particle. Red represents the particles binding to mitochondria and blue represents the particles not binding to mitochondria. n=34. (B, C) Simple logistic regression analysis of (A). This predicts a probability of mitochondrial binding of the Clu particle depending on the particle speed (B) or size (C). The value 1 of the y-axis represents mitochondrial binding of the particle, and 0 represents mitochondrial non-binding. The solid line indicates the mean of the probability, and the dotted line indicates a 95% confidence interval. A logistic Regression test was performed using GraphPad Prism.

### Supplementary movies

**S1 Movie**. **Live-imaging of the follicles from *clu^CA06604^*/+; *nos GAL4*/*UASp-FLClu*::*mScarlet* female.** Endogenous Clu GFPTrap (green) and ectopic FLClu::mScarlet (magenta) were recorded at 11-second intervals for 5 minutes using a Nikon A1 confocal laser scanning microscope. Video was recorded at ten frames per second.

**S2 Movie. Live-imaging of the follicles from *clu^CA06604^*/+; *nos GAL4*/*UASp-clu𝛥DUF****::**mScarlet*** **female.** Endogenous Clu GFPTrap (green) and ectopic 𝛥DUF::mScarlet (magenta) were recorded at 4.2-seccond intervals for 2 minutes using a Nikon A1 plus confocal microscope. Video was recorded at ten frames per second.

**S3 Movie. Live-imaging of the follicles from *clu^CA06604^*/+; *nos GAL4*/*UASp-clu𝛥Clu****::**mScarlet*** **female**. Endogenous Clu GFPTrap (green) and ectopic 𝛥Clu::mScarlet (magenta) were recorded at 2.1-second intervals for 2 minutes using a Nikon A1 plus confocal microscope. Video was recorded at ten frames per second.

**S4 Movie. Live-imaging of the follicles from *clu^CA06604^*/+; *nos GAL4*/*UASp-clu𝛥TPR*::*mScarlet* female.** Endogenous Clu GFPTrap (green) and ectopic 𝛥TPR::mScarlet (magenta) were recorded at 2.1-second intervals for 2 minutes using a Nikon A1 plus confocal microscope. Video was recorded at ten frames per second.

**S5 Movie. Live-imaging of the follicles from *clu^CA06604^*female exposed to puromycin.** Dissected ovarioles were exposed to 10 µM puromycin and recorded at 20-second intervals for 10 minutes using a Nikon Eclipse Ti2 spinning disk microscope. Video was recorded at ten frames per second.

**S6 Movie. Live-imaging of the follicles from well-fed *clu^CA06604^* female.** The object was recorded at 0.2-second intervals for 3 minutes using a Nikon Eclipse Ti2 spinning disk microscope. Video was recorded at ten frames per second.

**S7 Movie. Live-imaging of the follicles from well-fed *clu^CA06604^* female exposed to cycloheximide.** Dissected ovarioles were exposed to 3.5 mM cycloheximide for 20 minutes and recorded at 0.2-second intervals for 3 minutes using a Nikon Eclipse Ti2 spinning disk microscope. Video was recorded at ten frames per second.

**S8 Movie. Live-imaging of the follicles from starved *clu^CA06604^*female exposed to insulin.** Dissected ovarioles from starved female were subjected to 100 µg/mL insulin. The object was recorded at 20-second intervals for 20 minutes using a Nikon Eclipse Ti2 spinning disk microscope. Video was recorded at ten frames per second.

**S9 Movie. Live-imaging of the follicles from starved *clu^CA06604^*female exposed to CHX followed by insulin.** Dissected ovarioles from starved female were treated with CHX followed by 100 µg/mL insulin. The object was recorded at 20-second intervals for 20 minutes using a Nikon Eclipse Ti2 spinning disk microscope. Video was recorded at ten frames per second.

**S10 Movie. Live-imaging of the follicles from well-fed *clu^CA06604^* female exposed to hydrogen peroxide after dissection with insulin-containing CS media**. The object was recorded at 20-second intervals for 12 minutes using a Nikon Eclipse Ti2 spinning disk microscope. Video was recorded at ten frames per second.

**S11 Movie. Live-imaging of the follicle from well-fed *clu^CA06604^*female exposed to hydrogen peroxide following incubation with 3.5 mM CHX.** The object was recorded at 20-second intervals for 12 minutes using a Nikon Eclipse Ti2 spinning disk microscope. Video was recorded at ten frames per second.

**S12 Movie. Live-imaging of the follicle from well-fed *clu^CA06604^* female exposed to hydrogen peroxide following incubation with 7 mM CHX.** The object was recorded at 20-second intervals for 12 minutes using a Nikon Eclipse Ti2 spinning disk microscope. Video was recorded at ten frames per second.

**S13 Movie**. **Live-imaging of the follicles from Clu GFPTrap *clu^CA06604^* female labeled with 50 nM TMRE.** Clu GFPTrap (green) and mitochondria labeled with TMRE (magenta) were recorded at 4.3-second intervals for 10 minutes using a Nikon A1 confocal laser scanning microscope. Video was recorded at ten frames per second.

