## Supplemental Figure 1 and 2 for "*Drosophila* Clueless ribonucleoprotein particles display novel dynamics that rely on the availability of functional protein and polysome equilibrium"

### Supplementary Fig 1. Cycloheximide decreases Processing bodies *ex vivo*

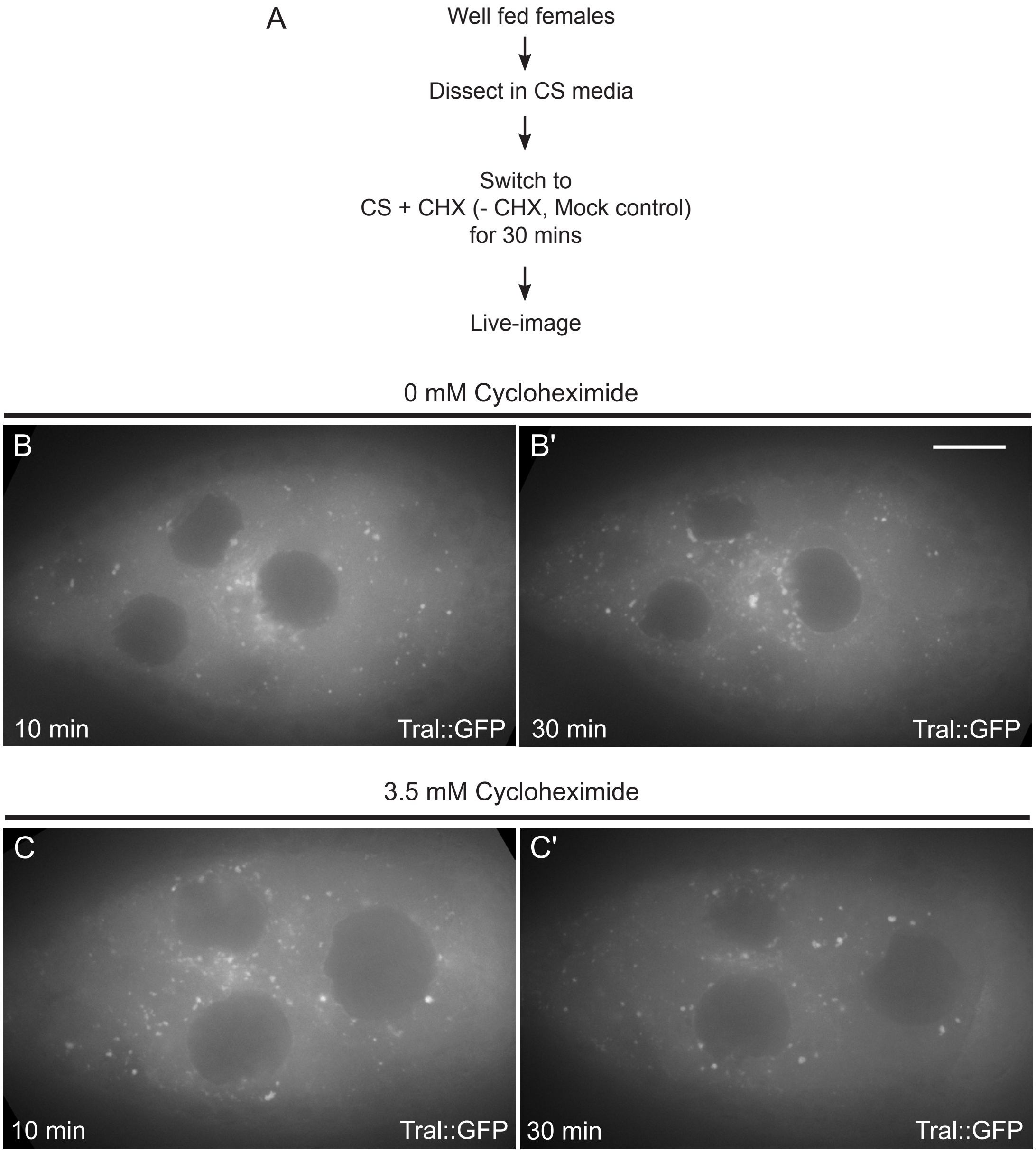

Supplementary Fig 2. Clu particles associated with mitochondria move more slowly

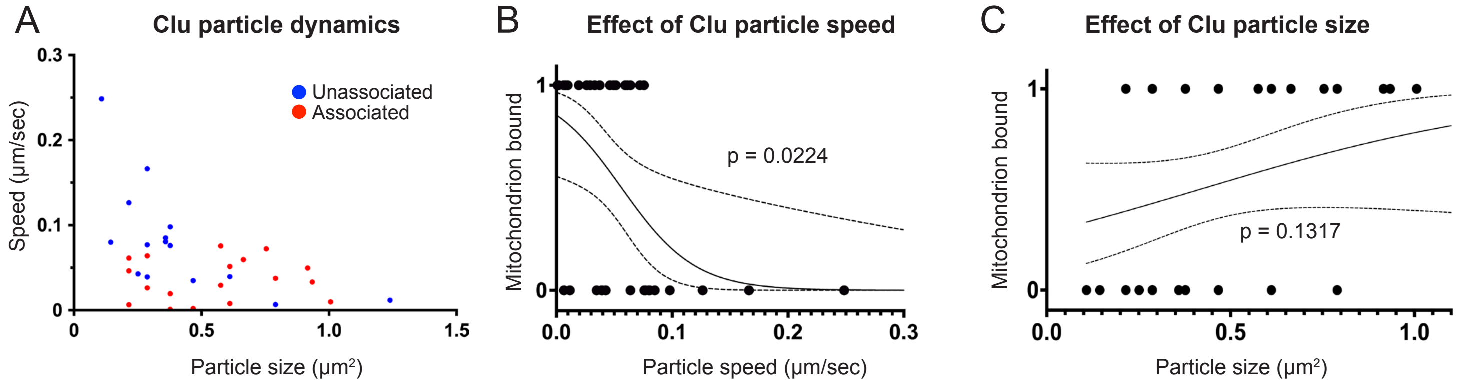
